# Clonal interference, genetic variation and the speed of evolution in structured populations

**DOI:** 10.1101/2025.02.22.639636

**Authors:** Yang Ping Kuo, Jiewen Hu, Oana Carja

## Abstract

When it comes to understanding the role that population structure plays in shaping rates of evolution, it is commonly accepted that interference between evolutionary innovations is more prevalent in structured populations compared to well-mixed, and that population structure reduces the rate of evolution, while simultaneously promoting maintenance of genetic variation. Prior models usually represent population structure using two or more connected demes or lattices with periodic boundary conditions. Fundamentally, the observed spatial evolutionary slow-down is rooted in the fact that these types of structures increase the time it takes for a selective sweep and therefore, increase the probability that multiple beneficial mutations will coexist and interfere. Here we show that systematically introducing more heterogeneity in population structure can reshape these prior conclusions and lead to a much wider range of observed evolutionary outcome, including increased rates of evolution. At a big picture level, our results showcase that the evolutionary effects of population structure crucially depend on the properties of the topology considered. Understanding these spatial properties is therefore requisite for making meaningful evolutionary comparisons across different topologies, for forming sensible null expectations about experimental or observational data or for developing an intuition for when well-mixed approximations, or approximations done using spatial models with a high degree of symmetry, might not apply.

## Introduction

The ability to efficiently accumulate beneficial mutations and innovations shapes a population’s adaptation and rate of evolution (Blokzijl et al., 2016; Bonnet et al., 2022; Markov et al., 2023). As beneficial mutants appear and rise in frequency in the population, competition between them can cause some of these evolutionary innovations to go extinct, impeding the evolutionary process. This is why interference between clones and its effects on rates of evolutionary dynamics has been the subject of substantial interest since the 1930s, when both Fisher (1930) and Muller (1932) first noted its potential importance (Fisher, 1930; Muller, 1932). In the well-mixed population limit, for a single asexual population assumed to climb a hill in fitness space, the amount of variation in fitness accumulating and competing within the population has been shown to be determined by a balance between selection, which destroys variation, and beneficial mutations, which create more. This balance depends on the population parameters: the population size, the beneficial mutation rate, and the distribution of the fitness increments of the mutants, with predicted adaptation speeds scaling with the logarithm of population size and mutation rate (Desai and Fisher, 2007; Rouzine et al., 2003; Brunet et al., 2008; Rouzine et al., 2008).

When it comes to understanding the role that population structure plays in shaping clonal interference and rates of evolution, it is commonly accepted that interference is more prevalent in structured populations compared to well-mixed, and that spatial or population structure reduces the rate of evolution, while simultaneously promoting maintenance of genetic variation (Martens and Hallatschek, 2011; Martens et al., 2011; Kryazhimskiy et al., 2012; Gordo and Campos, 2006). Prior models usually represent population structure using two or more connected demes or linear and hexagonal lattices with periodic boundary conditions (Martens and Hallatschek, 2011; Martens et al., 2011; Otwinowski and Krug, 2014). Fundamentally, the observed spatial evolutionary slow-down is rooted in the fact that these types of regular, symmetric spatial structures increase the time it takes for a beneficial mutant to sweep to fixation and therefore, increase the probability that multiple beneficial mutations will coexist, compete and interfere. This also allows the population to maintain genetic diversity for longer in the face of selection (Wakeley, 1998; Cherry and Wakeley, 2003; Schneider et al., 2016).

Here we ask if introducing more heterogeneity in the population structure considered can reshape these prior conclusions and lead to evolutionary outcomes not yet observed. As we move away from the limit of lattices and other isothermal structures (Lieberman et al., 2005; Kuo et al., 2024; Kuo and Carja, 2024a), with high levels of symmetry and regularity, do we still observe the spatial evolutionary slow-down, or are there structures that can, in fact, decrease interference and speed up rates of adaptation? Knowing how the effects of population structure depend on the properties of the topology considered is crucial for making meaningful comparisons across different structures (Poon et al., 2021) and for developing an intuition for when and why well-mixed approximations, or approximations and inferences done using models with a high degree of spatial symmetry, might not apply.

To represent more heterogeneous population topologies, we use networks as mathematical proxies for the structure of reproduction and replacement of a population and we systematically study how the accumulation of beneficial mutations depends on the graph’s properties. The nodes of the graph can be assumed to either constitute individuals in the population, with edges representing the local pattern of replacement and substitution, or subpopulations connected by edges as migration corridors. Networks make for a natural and versatile mathematical representation of heterogenous population structure because they allow for continuous tuning of the complexity of the population topology: the limit of a well-mixed population can be represented using the complete graph, other regular structures, such as lattices, can be represented using k-regular graphs and, through repeated and systematic deletion or addition of links between nodes, we have a lot of control over the topological properties we want to design and study.

Another benefit of using the mathematical proxy of networks is the existence of a previous body of theoretical work focused on the study of how graph structure shapes evolutionary dynamics (Lieberman et al., 2005; Antal et al., 2006; Frean et al., 2013; Monk et al., 2014; Pavlogiannis et al., 2017; Sharma and Traulsen, 2022; Svoboda et al., 2024; Kuo et al., 2024; Kuo and Carja, 2024b,a). Prior models have shown that the addition of heterogeneity to population structure can greatly extend the range of possible evolutionary outcome for a single new mutant entering the population. For example, under the assumptions of the standard Moran model, there exist graphs that can change the reproductive success and probabilities of fixation of a new mutant, in contrast to what had been observed using symmetric deme-based and lattice-based structures (Wright, 1943; Kimura and Weiss, 1964; Carja et al., 2014; Maruyama, 1970; Slatkin, 1981; Whitlock and Barton, 1997; Whitlock, 2003; Cherry and Wakeley, 2003) and graphs can be classified into three different categories: amplifiers, isothermal (a class which includes lattices and other regular structures) and suppressors topologies (Adlam et al., 2015; Hindersin and Traulsen, 2015; Allen et al., 2021; Tkadlec et al., 2020; Kuo et al., 2024; Kuo and Carja, 2024b,a). At a big picture level, this is because, heterogeneity in a node’s number of neighbors creates heterogeneity in node reproductive advantage, with some node individuals more likely to reproduce than others and some more likely to die (Kuo and Carja, 2024a; Kuo et al., 2024; Kuo and Carja, 2024b).

While network representations can be very versatile, they can also present significant challenges. One of the main obstacles to theoretical progress in evolutionary graph theory has been the analytical difficulty of multidimensional stochastic models, and most previous results have been restricted to studying times and probabilities of fixation for a single new mutation appearing on the graph (Hindersin and Traulsen, 2015; Allen et al., 2021; Tkadlec et al., 2020). These previous single-mutation frameworks can only be used to predict sequential fixation dynamics, in the limit of small mutation rates, when there are at most two variants present in the population. Due to the difficulty of obtaining closed form solutions, prior models usually also studied one network family at a time, making their conclusions seem unrealistic for most datasets and difficult to apply to inferred, previously unseen structures that might not conform to a pre-defined class (but see Kuo and Carja (2024a); Kuo et al. (2024); Kuo and Carja (2024b)).

Another challenge stems from the fact that the flexibility of networks can also make them mathematically ‘messy’, they can have tens of properties that can be quantified (from diameter, to clustering coefficient, to transitivity, degree distribution, amount of bottlenecks) and most of these properties are hard to study independently from one another. However, in previous work we have designed algorithms to control one network property at a time and have shown that graphs are not as difficult to study as they may seem. We have shown that we can make significant analytical progress by computing and understanding two main evolutionary properties of the networks: the amplification factor and the acceleration factor (Kuo and Carja, 2024a). The amplification factor essentially quantifies by how much the network topology can increase fixation probabilities of new beneficial mutants, compared to a well-mixed population, while the acceleration factor quantifies by how much the topology speeds up or slows down their spread through the population.

By designing a model which combines evolutionary graph theory and mutational interference theory, our results here provide new insights in each respective field. In agreement with previous lattice and deme-based models, our results show that degree-homogeneous network structures (k-regular graphs) reduce the rate of beneficial mutation accumulation, by slowing mutant fixation and allowing for other mutants to arise and interfere with the existing ones. However, in contrast with previous models, we show that, as we move away from the limit of k-regular graphs, more heterogenous topologies can reduce rates of interference and increase the rate of accumulation of beneficial mutations.

For example, accelerator graphs, by speeding up mutant fixation, can decrease rates of clonal interference (**Figure 1A**). Similarly, suppressor topologies, where beneficial mutants are less likely to survive stochastic drift and probabilities of fixation of beneficial mutations are reduced, can also reduce the probability that multiple established mutant lineages interfere with one another. We refer to this effect as the benefit of suppression. Conversely, one can think of this as the cost of amplification, since amplifier topologies, by increasing the spread of new beneficial mutants, can lead to higher rates of interference and slow down the evolutionary process (**Figure 1B**). When does this cost of amplification exceed their benefit of increased probabilities of beneficial mutant fixation? The answer is not straightforward, since most networks affect both probabilities and times to fixation and it is ultimately the interplay of both that shapes rates of evolution (Hindersin and Traulsen, 2014; Tkadlec et al., 2019; Kuo and Carja, 2024b,a).

**Figure 1:**
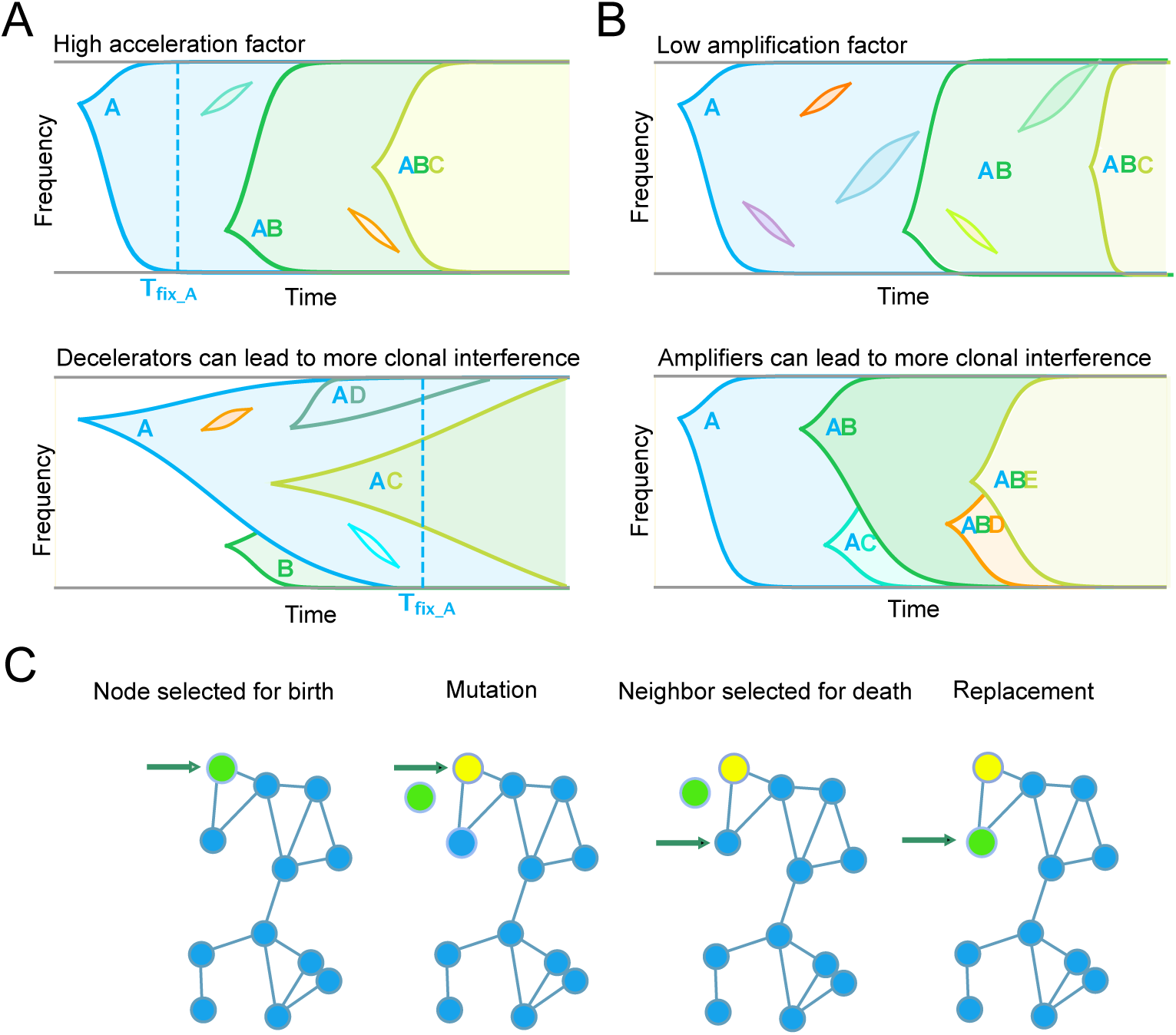
For beneficial mutations to accumulate, they must both arise and fix in a population. Heterogeneity in the structure of reproduction and replacement can affect both the probability and time of spread of new mutations in the population and can thus reshape clonal interference and rates of evolution. **Panel A.** Intuition on how topologies that decelerate mutant spread can increase rates of clonal interference. Two different scenarios are shown. Top: Assume an accelerator network. As beneficial mutations enter the population, their time to fixation is on average smaller than the waiting time for a new mutation to appear and mutations sweep sequentially, with little clonal interference. Bottom: For the same mutation rate, a population structure that increases the time to fixation of new beneficial mutants (decelerator structures), such that it becomes longer than the expected time of new mutants entering the population, can lead to more clonal interference. As a consequence, established beneficial mutations can be wasted (B mutants) unless they happen to occur on the background of the winning clone (C mutants). **Panel B.** Intuition on how topologies that amplify mutant spread can increase rates of clonal interference. Heterogeneous population topologies can change the establishment probability of beneficial mutants entering the population, leading to two drastically different adaptation scenarios. (Top) Lower establishment probability leads to a sequential fixation regime. (Bottom) For the same mutation rate, amplifier structures, which increase probabilities of mutant establishment and fixation, can lead to higher rates of clonal interference. **Panel C.** The Bd (Birth-death) update rules with constant supply of mutation. At each time step, in the Moran process, a node is selected for birth proportional to fitness (birth step, green node). The offspring replaces a randomly selected neighbor node (the death step). A mutation occurs with probability *U* and is assigned to the initial birth node (yellow node). The offspring of the initial node replaces the node selected for death.

To quantitatively answer these questions and rigorously understand the role of network structure in shaping mutation accumulation, we derive analytic expressions for the rate of evolution in terms of the two evolutionary network properties, amplification and acceleration, and show that these approximations are sufficient for obtaining a unified understanding of the role of heterogeneous population topology in shaping evolutionary dynamics. We also derive the fundamental theorem of natural selection that links the speed of evolution to genetic variation (Fisher, 1999) for population topologies of arbitrary heterogeneity.

Lastly, we discuss how to apply these network-based approaches to large-scale datasets of biological and social organization. We study the rate of evolution in two natural systems with drastically different spatial architectures and organization, a cellular and a social system. Our results show that both cellular and social architectures reduce the rate of innovation accumulation under biologically relevant mutation rates, compared to well-mixed populations. However, the two spatial structures show very different behavior when it comes to the mutational variation in the population. Cellular networks tend to have the same or lower mutation variation as the well-mixed population. In contrast, social networks promote maintenance of higher levels of cultural diversity. If we hypothesize that these networks are themselves shaped by natural selection, our results hint at a potential tug-of-war between optimizing for speed or variation in these systems. Overall, our results provide the foundation for further exploration of the role of heterogeneous spatial topology in shaping the dynamics of adaptation.

### Model

To study how heterogenous population structure shapes patterns of genetic variation and long-term rates of adaptation in the concurrent mutations regime, we study an asexual population of *N* haploid individuals. We use the infinite sites mutation model (Kimura, 1969), and assume that 1) there are *b* beneficial sites where mutations can occur, 2) *b* is assumed to be very large such that no two mutations occur at the same site, and 3) there is no recombination. New beneficial mutations appear at a rate *µ* per site and we study a model in which each beneficial mutation has the same effect, *s*, on fitness (i.e., each fitness step uphill is of the same size). The total beneficial mutation rate per replication is therefore *U* = *µb*. There are no epistatic interactions between mutations, such that fitness is multiplicative and an individual with *l* mutations has reproductive fitness of (1 + *s*)*^l^* ≈ 1 + *ls,* for *s* ≪ 1. Two mutants can have the same number of mutations, *l,* at different sites. For the purpose of this paper, we refer to them collectively as *l*-mutants, or in the *l*-mutant class. The interference process in our model is more closely related to the one in Desai and Fisher (2007) than to the one in Gerrish and Lenski (1998), since *s* is a constant and not drawn from a probability distribution. We focus on the effects of positive selection and neglect neutral or deleterious mutations, since even this simplest possible model with many equal-strength beneficial mutations is not understood and provides an initial first step towards developing a multi-mutation theory.

To represent heterogeneous population spatial structure, we use unweighted and undirected graphs, where each node in the graph represents one individual in the population and edges are proxies for the local pattern of reproduction and replacement. We can also interpret a node as a homogeneous subgroup of individuals and the edges as migration corridors between them, with the assumption that the timescale of a mutation traveling between nodes is much larger than the timescale of fixation within a node. We use a Moran Birth-death (Bd) process to track mutant frequency changes on the network (**Figure 1C**). The Birth-death process assumes that, at each time step, an individual from the population is selected to reproduce proportional to fitness and a random neighbor node is selected to die, leaving an unoccupied node for the offspring of the reproducing node. This is the essential difference that allows us to study the role of local population structure, in comparison to a well-mixed population: in a well-mixed population, a node from the entire network would be randomly selected for death. Each time an individual is selected to reproduce, with probability *U*, it can acquire an additional beneficial mutation and we make the arbitrary choice to assign the mutation to the initial reproducing node.

We study the role of population structure in shaping rates of evolution by analyzing how graph properties determine the speed at which the population accumulates beneficial mutations. We use graph generation algorithms implemented in NetworkX (Hagberg et al., 2008) and analyze a wide variety of graph families including preferential attachment graphs (Barabási and Albert, 1999), Erdős-Rényi graphs (Erdős and Rényi, 1960), bipartite graphs (Asratian et al., 1998), small world networks, used to model properties of social networks (Watts and Strogatz, 1998), and random geometric networks, mathematical representations of populations embedded in Euclidean space (Waxman, 1988; Penrose et al., 2003).

In order to design network topologies with pre-determined properties, that might not belong to these graph families, we use algorithms we have previously described that allow us to systematically tune network parameters independently of one another (Kuo et al., 2024; Kuo and Carja, 2024b). In previous work we have shown that, for single mutations, the evolutionary role of complex spatial structure can be quantified by analyzing two essential network properties: the network amplification factor, which shapes probabilities of fixation compared to well-mixed populations (Kuo et al., 2024; Kuo and Carja, 2024a) and the network acceleration factor, which shapes the time to fixation for new mutations in the population (Kuo and Carja, 2024a,b). The network amplification factor *α* quantifies how to rescale the selection coefficient *s* for the well-mixed model to obtain the same probability of fixation as an allele with selection coefficient *s* on a network population. In other words, a mutation with a selective benefit *s* in a graph population, would have a probability of fixation corresponding to an equivalent selection coefficient *αs* in a well-mixed population. If *α >* 1, the graph amplifies selection, if *α <* 1 the graph suppresses selection and if *α* = 1 the graph does not change probabilities of fixation compared to the well-mixed. Similarly, we previously introduced the network acceleration factor, *λ* (Kuo and Carja, 2024b), and showed that this network property shapes times to fixation, making the evolutionary dynamics *λ* times faster than in a well-mixed population, for *λ >* 1. The networks we use here allow us to continuously vary these two parameters and investigate their distinct roles in shaping evolutionary dynamics. We present a detailed list of the graphs used in this study, as well as details on the computation of graph properties studied in the **Materials and methods** section.

In addition to the two main evolutionary parameters of spatial structure, the three other important parameters in this model are the fitness increase provided by each mutation, *s*, the population size, *N* and the beneficial mutation rate per individual per generation, *U*. In what follows, we write analytic predictions for the speed and rate of evolution and the mutational variation as functions of these five parameters (as well as other statistical properties of the spatial structure) and compare these analytic predictions with results from Monte Carlo simulations. These analytic results allow us to generalize our insights beyond the graph families and simulations discussed here.

For the simulations, we initialize all nodes to be wild-type, with zero mutations. We run the simulation until steady state, where the loss of less fit clones due to drift and selection balances out the supply of new beneficial mutations that survive drift. We find that *N* (*N* −1) Birth-death steps after the first mutant appears is enough to achieve this, since *N* (*N* − 1) is the mean conditional extinction time of the wild-type under neutral selection (Ewens, 2004). After the simulation reaches the steady state, we set *t* = 0 and the lowest non-extinct fitness class as the new “0”-mutant, and we begin tracking the average number of mutations gained in the population. We also track the variance in the number of mutations in the population. We simulate 1000*/µ* steps after steady state is reached, which is the expected time for 1000 additional mutations to appear in the population. We run at least 1000 simulations for any given network structure and average the trajectories to obtain the mean and the variance of the number of mutations in the population. This ensures our results are averaged over at least ∼ 10^5^ mutations.

Figure 2A shows example average trajectories for the mean mutation count, accumulating linearly, after steady state has been reached. We use the slope of the trajectory as a measure of how fast the population is accumulating mutations and define this as the speed of evolution, *V*. We define the normalized rate of evolution, *v*, as (*NU*)*^−^*^1^*V*. While the speed of evolution *V* informs on how fast a population accumulates mutations, it does not directly inform whether interference between mutations is occurring. The rate of evolution *v*, on the other hand, quantifies, on average, by how much every mutation entering the population increases the number of accumulated mutations in the population. Since time is measured in units of generations divided by *NU*, in Figure 2A the slope of the average mutational trajectory is equivalent to *v*. In the limit of weak mutation, *v* represents the probability of fixation. As the mutation rate increases, *v* becomes smaller than the probability of fixation, since multiple lineages that are supposed to fix collide with one another, resulting in the loss of mutations, as seen in the decrease in slope with increasing mutation rate (Figure 2A). The mutational variance *σ*^2^ is determined by taking the variance in mutation number at every time step and averaging across all time steps and simulations.

**Figure 2:**
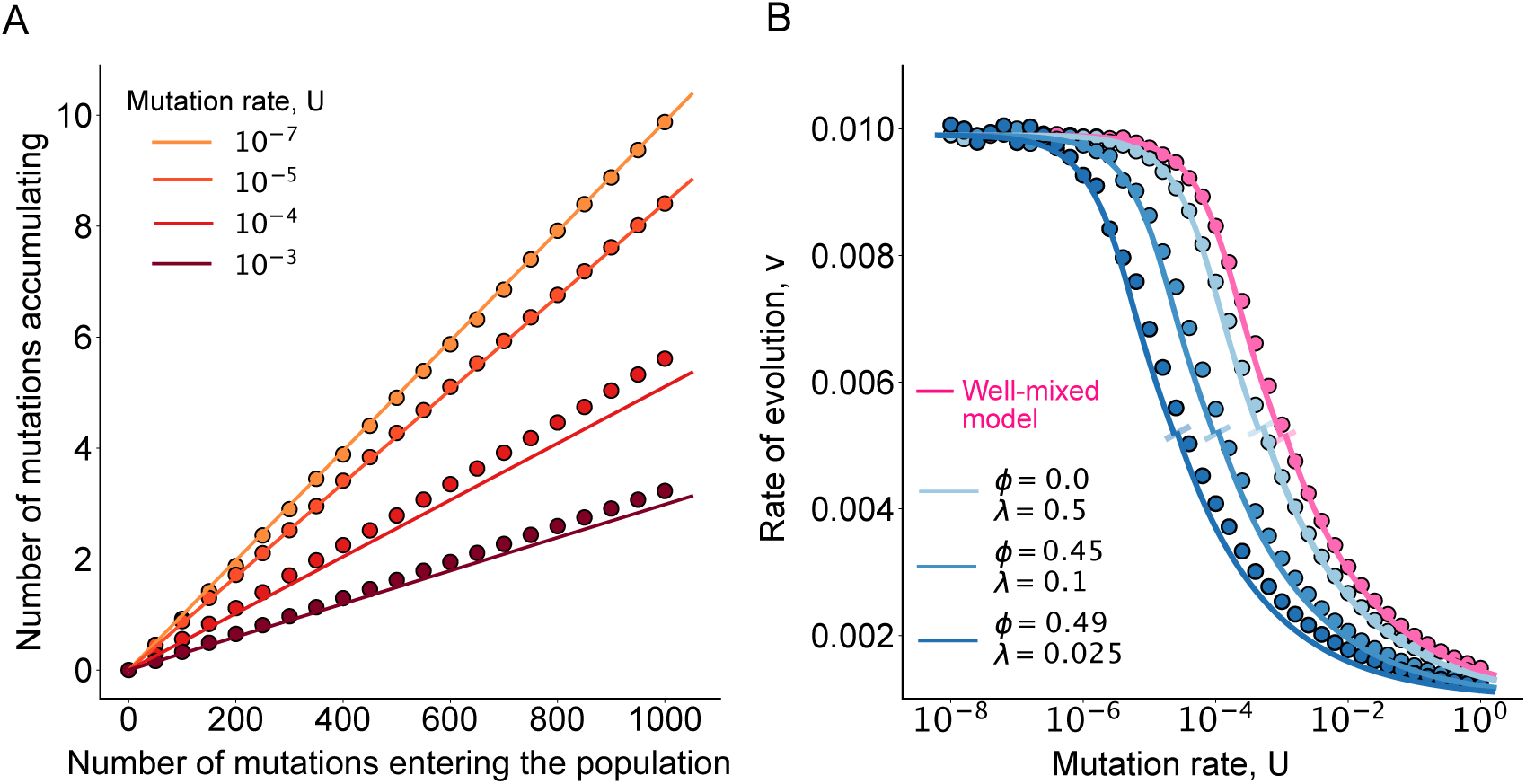
Rates of evolution for homogeneous population structures. Here dots represent rate of evolution calculated from ensemble averages across at least 10^3^ replicate Monte Carlo simulations per parameter, per network. The solid colored lines represent the theoretical rate of evolution for the network-structured population, calculated using equation (9). Short lines at 90*^◦^* angle highlight the transition between the two approximation regimes. **Panel A.** Average trajectories of the mean number of mutations accumulating in the population under different mutation rates. Time is normalized by dividing by *NU*. The population structure used is a 3-regular graph (with amplification factor equal to 1). Here *N* = 1000, *s* = 0.01 and *λ* = 0.1. **Panel B.** Rate of evolution, *v*, as a function of the mutation rate. Here *N* = 1000, *s* = 0.01 and the networks used are 3-regular graphs with amplification *α* = 1 and acceleration *λ* and number of cliques *ϕ* as depicted. As mutation rate increases, graphs with higher acceleration factor, by speeding up mutant establishment and spread, enter the clonal interference regime later and have higher rates of evolution.

## Results

We first outline the key analytical steps of our approach and show that the effect of network structure on the dynamics of mutation accumulation can be rigorously described using just the two main network evolutionary properties: the network amplification factor, *α*, and network acceleration factor, *λ*. The full analytical derivation is described in the **Supplementary Material**. In the following sections, we make comparisons between our analytic results and simulations and discuss the roles of each of these two network parameters independently, as well as evolutionary dynamics uniquely resulting from their interplay.

### Intuitive heuristics and outline of analytical derivations

Our analytical approximations rely on the diffusion approximation (Crow et al., 1970; Gardiner, 2009). Due to the added complexity of the network pattern of replication and replacement, direct application of diffusion can be mathematically very difficult. Prior analytic approaches developed for evolutionary dynamics of single mutants have used the adjacency matrix of the network (which uniquely identifies the graph) to track transition probabilities in mutant frequency (Hindersin et al., 2016). These approaches however become intractable for even small network sizes, since they would involve analysis of a stochastic process with *b^N^* states, where *b* is the number of sites with beneficial mutations.

The approach we take here, and the core idea behind our approximation, is that we can greatly reduce the complexity of the problem by representing the network using its node degree distribution. We track mutant frequencies across groups of nodes of the same degree and essentially transform the problem into a finite island population type model. While the degree distribution might not uniquely represent the network and some of the graph information is lost, this approach nonetheless shows great fit to the simulation results and, importantly, greatly reduces the number of possible states in the Moran model, making it applicable for networks of thousands of nodes and high degree of heterogeneity.

Our analysis is split into two parts. In the first part (**Supplementary Material Sections 1.1** and **1.2**), we first show that if we assume nodes of the same degree in the network to be “evolutionarily” identical and group them together, we can derive the expected change in mutant frequencies, for each group of nodes. To this end, we first observe that, since wild-type to mutant replacements depend on the number of edges between mutants and wild-type individuals, we can write this expected change in mutant frequencies as a function of the frequencies of the different edge types possible in the graph. After deriving the expected change in edge-type frequencies, we show that, if we assume weak selection and weak mutation, the *l*-mutant frequencies for each group of nodes with distinct degree evolve to the same quasi-equilibrium frequency. We define this quasi-equilibrium frequency as *x_l_*, which depends on the initial *l*-mutant placements on the network. The quasi-equilibrium edge frequencies also depend on *x_l_*. Selection and mutation, however, will slowly alter this quasi-equilibrium.

In the second part of the analysis (**Supplementary Material Sections 1.2** to **1.6**), we write the stochastic differential equation describing the evolution of the mutant frequencies. To do so, we replace the quasi-equilibrium *l*-mutant frequencies *x_l_* with their stochastic counterparts, random variables *X_l,t_.* The collection of random variables **X_t_** = *X*_0_, · · · *, X_l_,* · · · *, X_b_* evolves according to

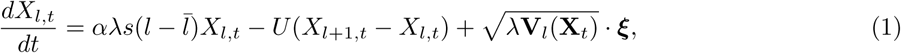

where *l̄* is the mean number of mutations in the population, **V***_l_* = *V_l_*_0_, · · · *, V_lm_* is the correlation in frequency change with other mutants, and ***ξ*** is a multivariate Gaussian noise variable.

In the limit of weak mutation, when there are at most two fitness classes present in the population, equation (1) recaptures results in evolutionary graph theory with one spreading mutant, showcasing the independent effects of the two network parameters *α* and *λ*. To show this, let us assume the fitter mutant class has frequency *X_t_* and the less fit class (1 − *X_t_*). Equation (1) reduces to

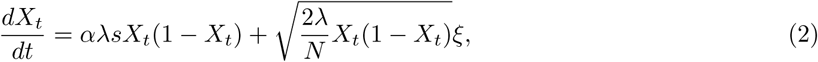

where *ξ* is Gaussian noise. Solving the Kolmogorov backward equation (Crow et al., 1970) yields the probability of beneficial mutant fixation

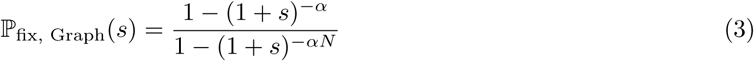

and, for weak selection, the conditional mean time to fixation can be written as

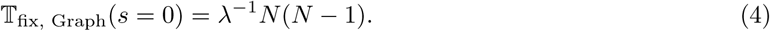

Equation (3) recaptures *α* as the amplification parameter, with the mutant on a graph *G* having the same fixation probability as a mutant with selection coefficient *αs* in a well-mixed population. Similarly, the graph accelerates the conditional mean time of mutant fixation by a factor of *λ*, compared to a well-mixed population. While we have previously shown that the amplification and acceleration factors have closed-form approximations using network descriptors, such as the mean and variance in degree (Kuo et al., 2024; Kuo and Carja, 2024b), they can also be computed using simulations: 1) estimate the probability of fixation and solve for *α* in equation (3) and 2) estimate the time to fixation and solve for *λ* in equation (4).

It is important to notice that equation (1) is similar to the one governing the equivalent Moran process for a well-mixed population, with appropriate parameter mapping changes that account for the roles of the two network evolutionary parameters, *α* and *λ*. In other words, if we set *α* = 1 and *λ* = 1, equation (1) has been widely studied to understand evolution in the well-mixed population limit (Tsimring et al., 1996; Cohen et al., 2005; Desai and Fisher, 2007; Rouzine et al., 2003).

This tells us that, in order to understand the evolutionary role of the network structure, we can use prior well-mixed approximations, with the right parameter ‘stretches’ that account for the evolutionary properties of the network. Even in this limit, a closed-form analytic solution has been notoriously hard to derive: the number of variables *b* is large, dropping the stochastic term leads to an infinitely large speed of evolution (Desai and Fisher, 2007), and, in general, moment equations do not close (Hallatschek, 2011).

The standard approach to find an approximation for the well-mixed model is to split the behavior of the mutant classes into two: 1) the “bulk”, where the mutant frequency is high and the drift term of equation (1) can be ignored and 2) the “front”, where the fitness class consists of the fittest mutants just entering the population, subject to the force of stochastic drift. The cutoff between the two classes, *q*, depends on a cutoff frequency *x_q_,* below which drift becomes important. Effectively, the difference between different prior analytical approaches is in defining where this transition occurs. Here, extra care needs to be taken to account for the population size of interest. For example, many of the assumptions in Desai and Fisher (2007) hinge on the population size being very large. We use graphs of sizes 1000-15000, which are large compared to previous evolutionary graph theory work, but small compared to the microbial populations Desai and Fisher (2007) are interested in.

For small mutation rate *U* (and small *NU* in our case), we use the criterion from Desai and Fisher (2007) and define the cutoff frequency *x_q_* = 1*/qs.* Following a similar derivation as theirs, we find the cutoff *q* to be

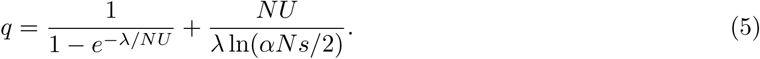

Using this *q*, we find the resulting speed of evolution to be

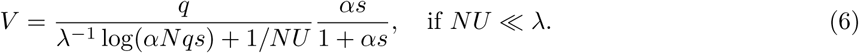

To obtain an approximation for large mutation rates (large *NU*), we use the criterion in Tsimring et al. (1996) and Cohen et al. (2005). Since, in this scenario, the mutation rate is assumed to be much higher than *s,* the mutant frequency at the stochastic edge, *X_q_*_+1_, depends solely on the *q*-mutant supplying mutations, rather than selection. Therefore, the cutoff frequency is independent of *s* and the choice of *x_q_* is often arbitrary and is selected to fit simulations. For example, Cohen et al. (2005) used *x_q_* = 4*/N.* Here, we find the speed of evolution

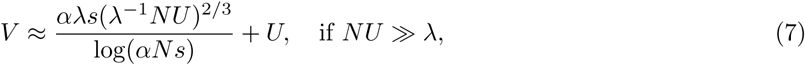

following the *U* ^2^*^/^*^3^ scaling first observed by Cohen et al. (2005) and Tsimring et al. (1996).

Once we write out the speed of evolution, we can easily approximate the amount of genetic variation in the population. We simply multiply equation (1) by *l*, sum over all *l,* and drop the stochastic drift term. This leads to the following relationship between *V* and *σ*^2^

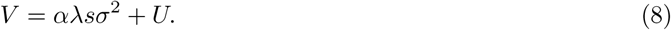

The stochastic terms can be dropped since drift is only prevalent when the mutant frequency is low, thus contributing very little to the variance, even when it drives the mutation class to extinction. More precisely, *x_l_* follows a Gaussian profile (Rouzine et al., 2003), so the force of drift matters at the tail, but *l*^2^*x_l_* falls off exponentially.

Equation (8) mirrors the fundamental theorem of natural selection for a well-mixed population, which states that the rate of fitness increase is equal to the variance in fitness: *sV* = *s*^2^*σ*^2^ (Fisher, 1999). The effect of the network structure is therefore represented by the coefficient *αλ* in front of *sσ*^2^. The extra factor, *U* is the result of recurrent mutation in our model. Equation (8) thus extends the fundamental theorem of natural selection for structured populations.

Using equations (6) and (7), we can now approximate *v* as

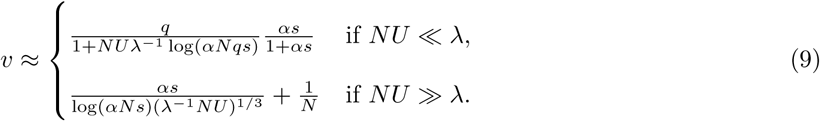

We show comparisons between these analytic approximations and simulations in Figure 2B and throughout the following figures. It is important to note here that the transition between the two forms of the approximation, driven by two different regimes for *NU*, and depending on network acceleration factor *λ*, can usually be easily observed in our figures and is denoted by an angled line (see, for example, the dark blue oblique line crossing the analytic approximation for *λ* = 0.025 around mutation rate *U* = 10*^−^*^4^ in Figure 2B).

### Degree-homogeneous structures and the role of the network acceleration factor

#### By decreasing clonal interference, graphs with increased acceleration factor increase rates of evolution

In this section, we use degree-homogenous isothermal graphs (amplification *α* = 1) to discuss the independent role the acceleration factor of a structure in shaping rates of evolution and connect our analytic results to other better-known network properties, such as average degree and higher-order network motifs. To do so, we use random *k*-regular graphs, where every node has exactly *k* connecting edges (lattice structures are a subset of this graph family). These types of graphs are known decelerators (Kuo and Carja, 2024b), with acceleration factor approximately equal to

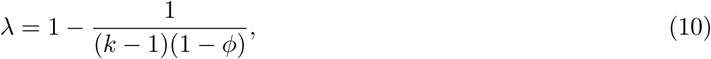

where *k* is the mean degree and *ϕ* is the fraction of cliques (cycles of length 3) in the network. In order to study the role of the acceleration parameter *λ*, we use degree-preserving edge swaps to tune this parameter, while keeping mean degree *k* constant (Kuo and Carja, 2024b).

We show the rate of evolution *v* as a function of the mutation rate *U* and the network acceleration factor *λ* in Figure 2B. Notice how mutation rate *U* and acceleration *λ* always appear together in equation (9). This means that the effect of the population structure can be thought of as creating an effective mutation rate *λ^−^*^1^*U.* In other words, a graph population would have the same rate of evolution *v* as a well-mixed population with mutation rate *λ^−^*^1^*U.* These homogeneous decelerator population structures effectively “shift” the rate of evolution curve to the left by ln(*λ^−^*^1^) and utilize beneficial mutations identically to a well-mixed population with a higher effective mutation rate. This explains the evolutionary slowdown previously observed for these types of topologies (Martens and Hallatschek, 2011; Martens et al., 2011; Kryazhimskiy et al., 2012; Gordo and Campos, 2006). Both decreasing the mean degree or increasing the fraction of cycles, i.e. decreasing the network acceleration factor, causes the time to mutant fixation/extinction to increase and this leads to an decrease in the rate of evolution (Figure 3). Even though *k*-regular graphs cannot have acceleration above the well-mixed structure (*λ <* 1), this result also points to the fact that it is possible for the rare of evolution to be increased by graphs with *λ >* 1.

**Figure 3:**
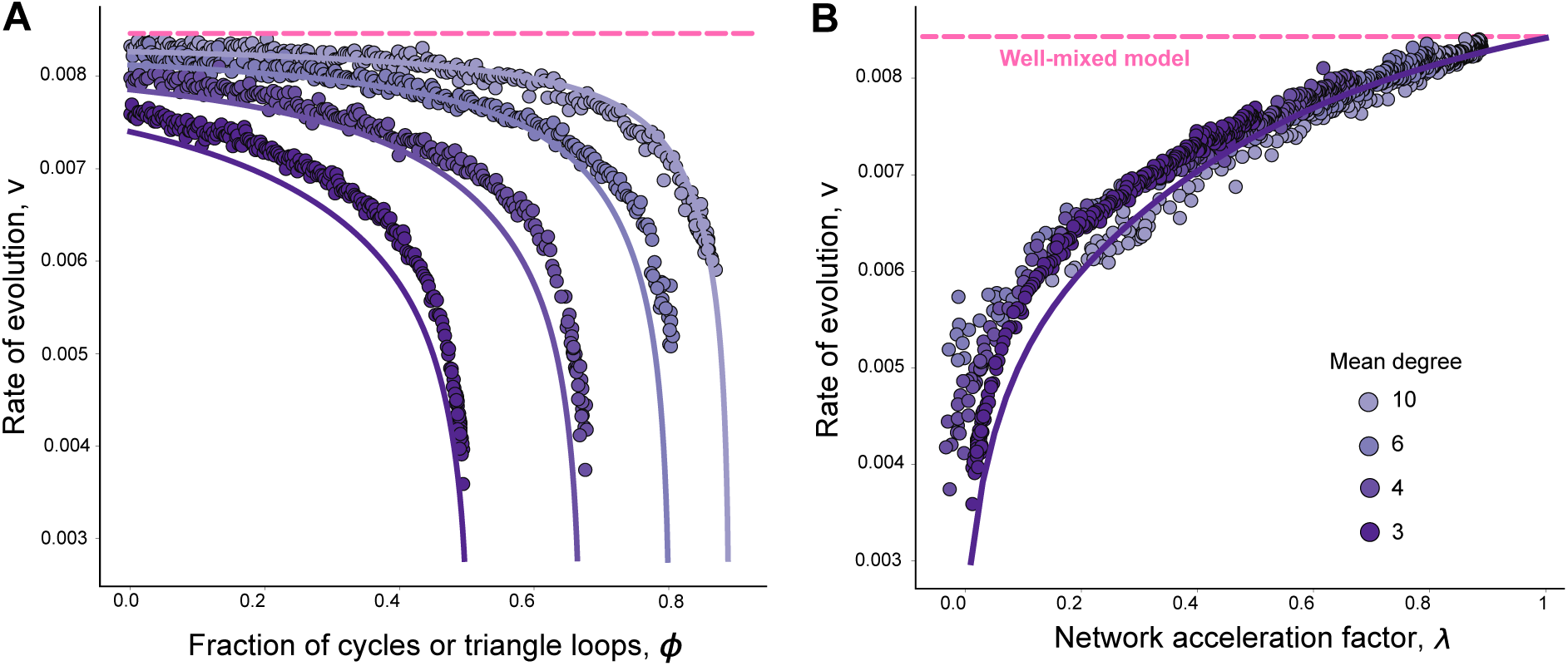
The role of mean degree and number of cycles (triangles) in shaping rates of evolution, for degree-homogeneous graphs. Here dots represent rate of evolution calculated from ensemble averages of at least 10^3^ replicate Monte Carlo simulations, per parameter, per network. The solid colored lines represent the theoretical rate of evolution for the network-structured population, calculated using equations (9) and (10). **Panel A.** Here, for a given mean degree, we vary the fraction of triangles in the graphs using previously described edge-swap algorithms. All graphs of the same color have the same mean degree, as in the legend, but different number of triangles, as on the *x*-axis. Graphs with high number of triangles decrease the network acceleration factor and thus decrease rates of evolution. Here, *N* = 1000, *s* = 0.01, *U* = 10*^−^*^4^. **Panel B.** We use the same graphs and parameters as in Panel A. We show that rates of evolution increase with network acceleration factor.

Since cycles in *k*-regular graphs shape times to fixation and thus the competition of mutant lineages, we visualize the role of these closed loops in Figure 4. Here all three graphs are 8-regular graphs, where every node has exactly 8 neighbors. The difference between the three networks lies in the fraction of closed triangle loops and, for a given mutation rate and selection coefficient, this difference can result in different evolutionary dynamics. The visualization can help provide an intuition into why these higher-order motifs change evolutionary patterns and lead to different mutation accumulation dynamics: since the spread of a mutant lineage critically depends on the number of edges that connect the mutant to other lineages (Kuo and Carja, 2024b), when the fraction of triangles is high, two nodes connected by an edge are likely to have shared neighbors. This leads to tightly connected clusters of nodes, where different clusters are connected by very few edges (Figure 4C). When a mutant lineage is spreading through the network, the lineage quickly takes over a highly interconnected cluster. The spread soon comes to a halt because the number of heterogeneous edges is consumed, as they only occur at the boundary connecting different mutant clusters. As the number of available heterogeneous edges decreases with the fraction of triangles, this decrease leads to higher times to fixation for new variants in the population.

**Figure 4:**
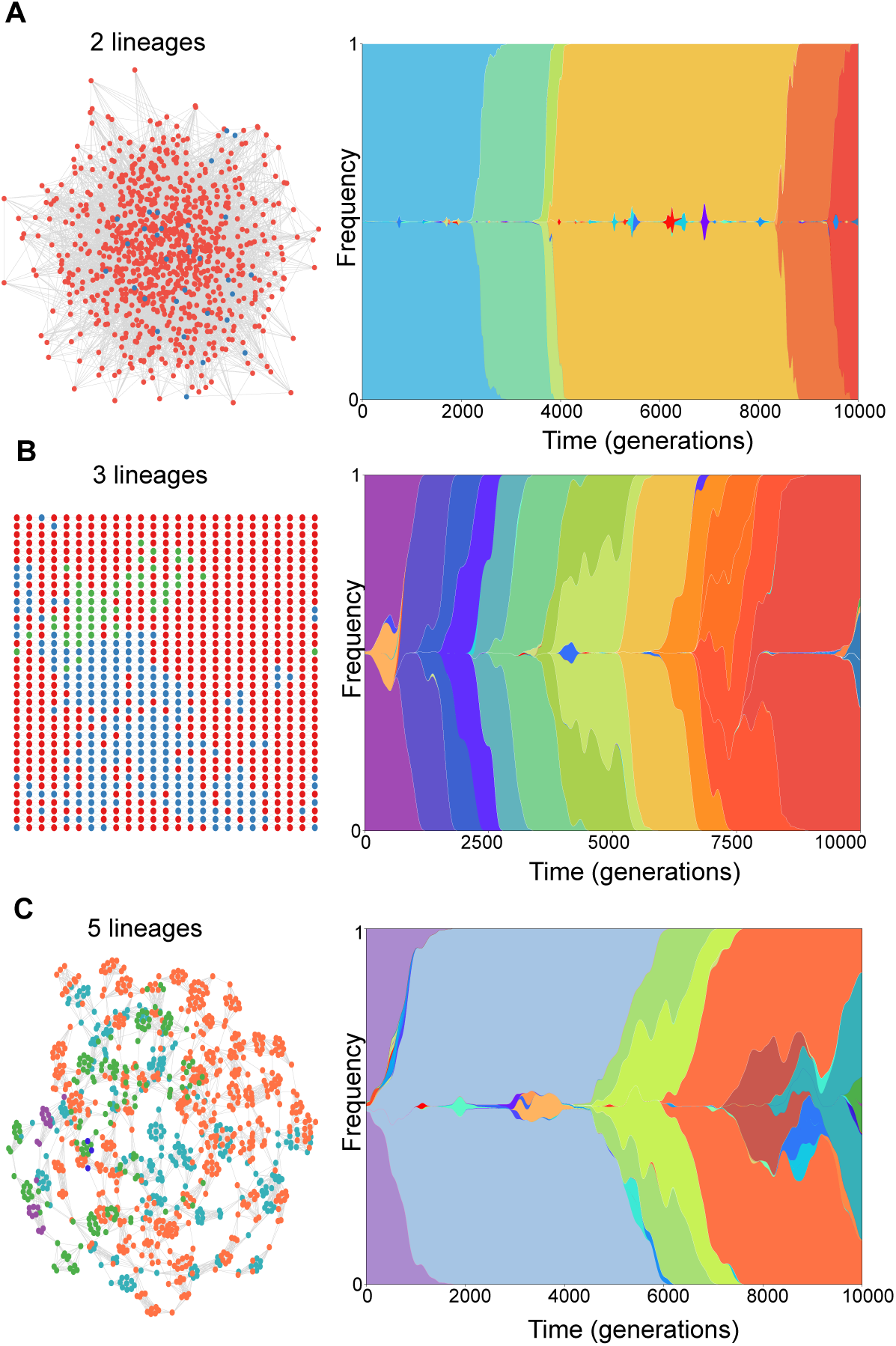
Intuitive visualization of the role cycles and closed loops play in decreasing acceleration factors and rates of evolution for degree-homogeneous graphs. Here the degree distribution is held constant (*k* = 8), as we vary the fraction of triangles in the graphs, using edge swapping operations. Population size *N* = 1000, *s* = 0.01, and *U* = 10*^−^*^4^. Fraction of triangles is 0 (top panel A), 0.428 (to showcase a lattice, in the middle panel B), and 0.85 (bottom panel C). Graphs with low triangles have higher acceleration factor leading to less interference. Left panels show distribution of mutants on different graphs after evolving for 10000 generations. Right panels show temporal evolution of clones in each populations.

#### Population structures with increased heterogeneity can reduce clonal interference and increase rates of evolution

Let us now focus on understanding the role of the amplification factor in shaping clonal interference and rates of evolution. To this end, we use graphs where we keep the acceleration factor constant (*λ* = 1, we show two example structures used in Figure 5A and B) and ask how introducing reproductive bias in the population structure affects the interference of advantageous mutations. Since the probability of fixation is tied to two aspects of mutation accumulation, i.e. how fast mutants fix and how likely two mutants are to compete, we cannot quantify how much interference occurs in these graphs by looking at the rate of evolution alone.

**Figure 5:**
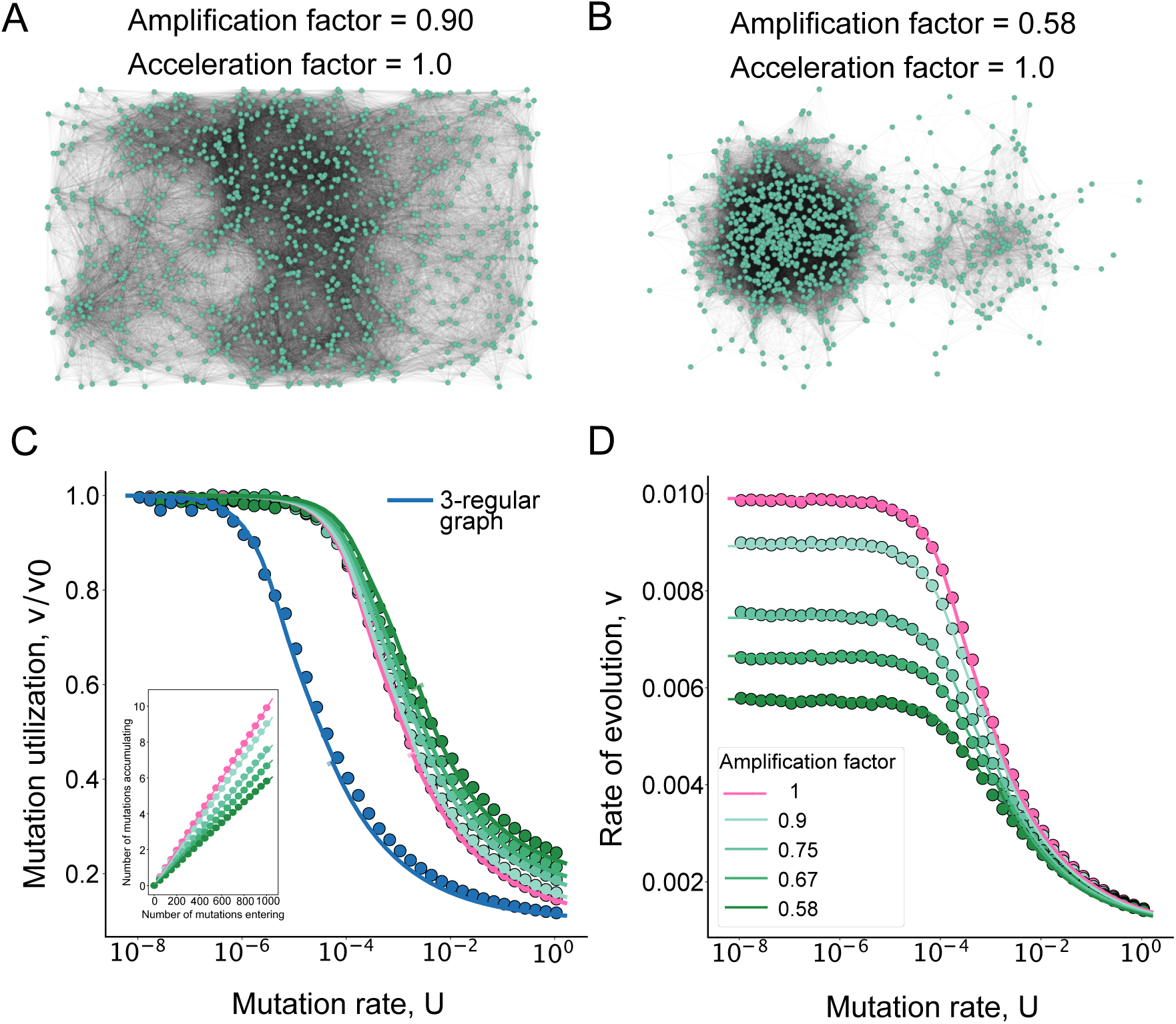
Spatial structures can reduce clonal interference compared to well-mixed or lattice-like populations. Panel. **A.** Spatial network with *α* = 0.9 and *λ* = 1.0. **Panel B.** Spatial network with *α* = 0.58 and *λ* = 1.0. **Panel C.** Mutation utilization is calculated by taking the rate of evolution, divided by the rate of evolution in the limit of vanishingly weak mutation rate. The dots are results from 1000 simulations per parameter, per network, divided by values from simulations of single mutation fixation for the same graph, while the lines are our analytical approximations. Short oblique lines highlight the transition between the two approximation regimes. Results for the 3-regular graph with *ϕ* = 0.49 from Figure 2 are included as comparison. The insert depicts mutation accumulation for the same structures. All networks have acceleration factor *λ* = 1 and amplification factor *α* as in the legend in Panel D. Population size *N* = 1000 and *s* = 0.01. **Panel D.** The lines represent analytical approximations using equation (9), while the dots are results from 1000 simulations per parameter, per network. Here, *N* = 1000 and *s* = 0.01. All networks have acceleration factor *λ* = 1 and amplification factor *α* as in the legend.

To specifically study clonal interference, let us define the rate of mutation utilization as *v/v*_0_, where *v*_0_ is *v* in the limit of weak mutation, equivalent to the probability of mutant fixation. For weak mutation rates, every mutation that’s destined to fix increases the mean number of mutations in the population by one. Therefore, in a sense, the mutation is being used with 100% efficiency. As the mutation rate increases, multiple lineages co-occur in the population, and multiple mutations that would otherwise be destined to fix are therefore needed to eventually increase the mean number of mutations in the population by one. Mutations are thus being used with lower efficiency, and many are wasted.

The decrease in mutation utilization as a function of mutation rate *U* is illustrated in Figure 5C. We show that graphs with a lower amplification factor utilize mutations more efficiently and thus reduce clonal interference. For comparison, we also include a degree-homogeneous structure (like the ones used in Figure 2) which shows the lower mutation utilization and the increased interference. Therefore, as the heterogeneity in the structure considered is increased, we can observe how space can reduce clonal interference, an outcome not expected for symmetric homogeneous structures and predicted by previous modeling approaches.

However, for these graphs with constant acceleration *λ* = 1, this cost of increased amplification (smaller rates of mutation utilization and higher rates of interference) does not outweigh the benefit of having a higher establishment probability, and the rate of evolution increases with amplification factor, with this difference diminishing as the mutation rate increases (Figure 5D).

#### Trade-offs between amplification and deceleration

While the previous analyses help us understand the independent roles of the two evolutionary parameters of spatial structure in shaping clonal interference and rates of evolution, most structures affect both the probability and the time to fixation for new mutants entering the population (i.e. *α* and *λ* are not 1). To conveniently study the interplay of acceleration and amplification, we use bipartite structures, which allow for an easy and intuitive analytical relationship between degree heterogeneity and the two network parameters, *α* and *λ*. This relationship is given by

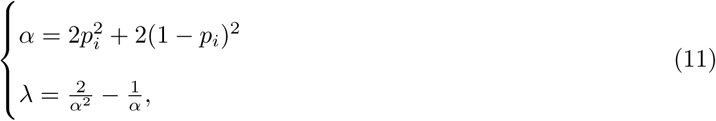

where *p_i_* and 1 − *p_i_* are the fractions of nodes in each of the two sets of the bipartite structure (see description in **Materials and methods**).

To obtain intuition in this regime, it helps to understand how the mutation rate can reshape the dynamics of mutation accumulation on the graph structure. In the limit of weak mutation (*U* = 10*^−^*^8^), sequential fixation dominates, and the rate of evolution is equal to the fixation probability of a single mutation. Since in this regime the time to fixation is less of a contributor, strong amplifiers of selection (usually structures with high degree heterogeneity) can accumulate mutations faster than other structures (Figure 6A).

**Figure 6:**
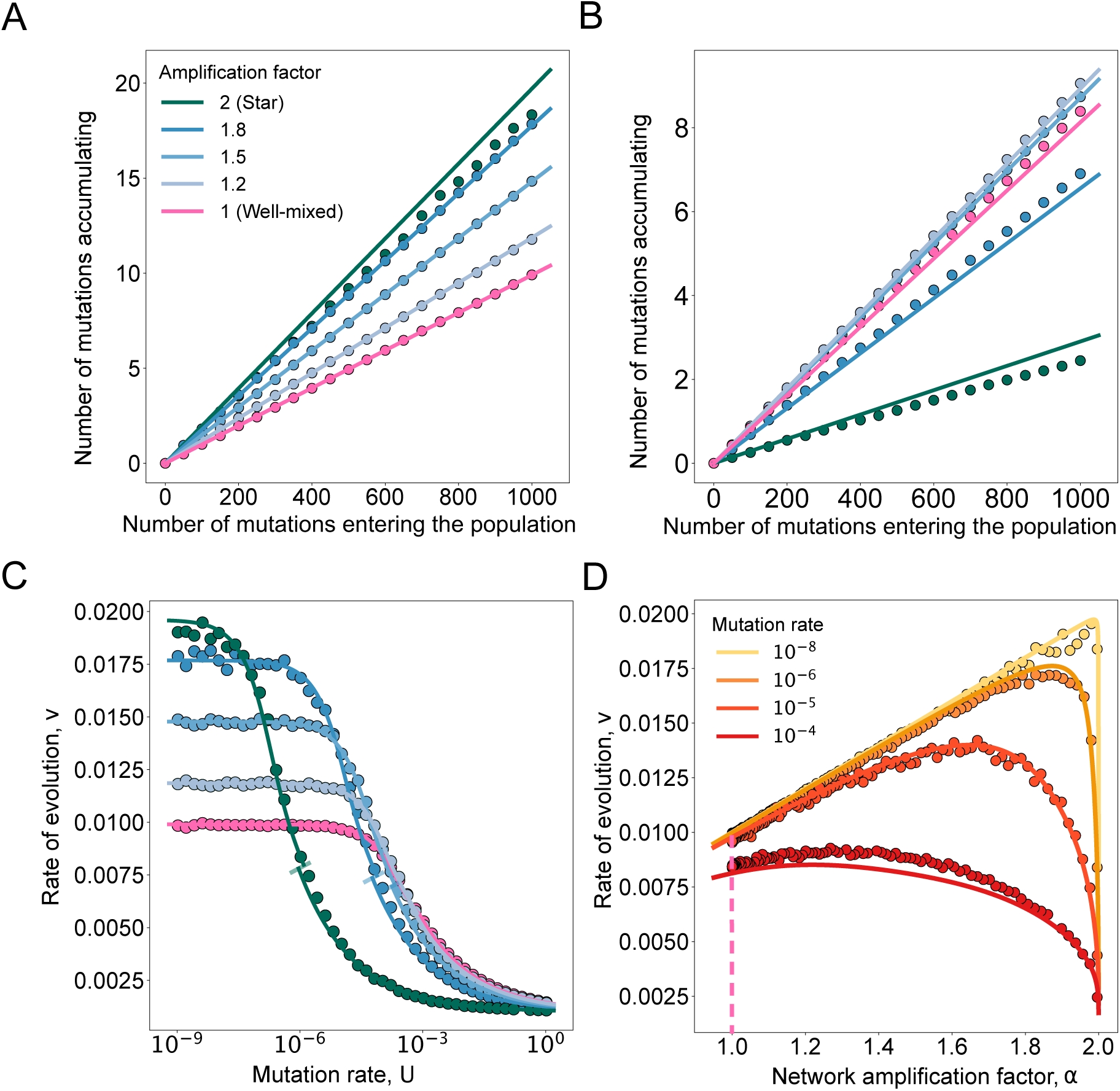
The mutation rate determines tradeoff effects between structure amplification and acceleration. The dots represent ensemble averages across at least 10^3^ replicate Monte Carlo simulations per parameter set and network. Here we use bipartite graphs with *α* and *λ* obtained from equation (11). Solid lines represent equation (9). Short oblique lines highlight the transition between the two approximation regimes. **Panel A.** Population size *N* = 1000, *s* = 0.01 and *U* = 10*^−^*^8^. **Panel B.** Population size *N* = 1000, *s* = 0.01 and *U* = 10*^−^*^4^. **Panel C.** Rate of evolution as a function of mutation rate. Same parameters as **Panel B**. **Panel D.** Rate of evolution as a function of the network amplification parameter.

As *NU* increases (*U* = 10*^−^*^4^), the rate of evolution depends not only on the probability of establishment, but also on the expected time for the mutants to fix and tradeoffs between amplification and deceleration can lead to strong amplifiers (like the star graph, with amplification factor of 2, acceleration factor of 0.001) having the lowest rates of evolution (Figure 6B). It is easy to understand the reason for this reversal. We saw in the previous section that, without taking deceleration into account (fixing *λ* = 1), the cost of increased amplification does not outweigh the benefit of having a higher establishment probability, and structures with higher amplification, *α*, have increased rates of evolution. However, for graphs where a higher probability of fixation also results in an increased time to fixation, the combined interference resulting from the cost of amplification and the increased time to fixation, can now outweigh the benefit of amplification. The amplifier structures where the total interference cost is larger than the benefit of amplification decrease rates of evolution. This result means that, contrary to single mutation evolutionary theory, weak amplifiers can exhibit the highest rates of evolution (Figure 6B).

We highlight this tradeoff between deceleration and amplification in Figure 6C. As the mutation rate increases, interference starts to become a dominant contributor to the rate of evolution, and amplifier networks can begin to act effectively as suppressors, decreasing the probability of mutant accumulation, compared to the well-mixed model (Figure 6C). For a mutation rate as low as 10*^−^*^8^, for example, the star graph begins to experience clonal interference and the rate of evolution begins to decrease. For larger population sizes, the star graph can experience clonal interference at even lower mutation rates, since *λ* ∼ 1*/N* decreases as population size increases. Figure 6D shows that even for an extremely weak mutation rate of *U* = 10*^−^*^8^, weaker amplifiers evolve faster than a star graph. This demonstrates that certain assumptions in single mutation evolutionary theory only hold at a knife’s edge — the probability of fixation is predictive of mutation accumulation only at the limit of really small mutation rates. Figure 6D also highlights that spatial structure can overall increase rates of evolution compared to well-mixed or lattice-like populations, for a wide range of mutation rates. We also explore more network families and show this result holds more generally in **Supplementary Figure S1**.

#### The fundamental theorem of natural selection on graphs: the relationship between rate of evolution and genetic variation

Our results provide a unifying way of predicting mutation accumulation by distilling properties of complex network organization through two single parameters: the acceleration and amplification factors. We compare the main families of networks in Figure 7A, under a mutation rate where the well-mixed graph begins to experience interference (*U* = 10*^−^*^4^). For this mutation rate, bipartite graphs have the highest rates of evolution out of all the graphs we explored, overtaking strong amplifiers like the star graph. They can also span a wide range of rates of evolution, showcasing that a network that is Pareto optimal for a single mutation in terms of fixation time and probability does not guarantee the best performance under an elevated mutation rate. The best-performing graph is inherently dependent on the mutational regime.

**Figure 7:**
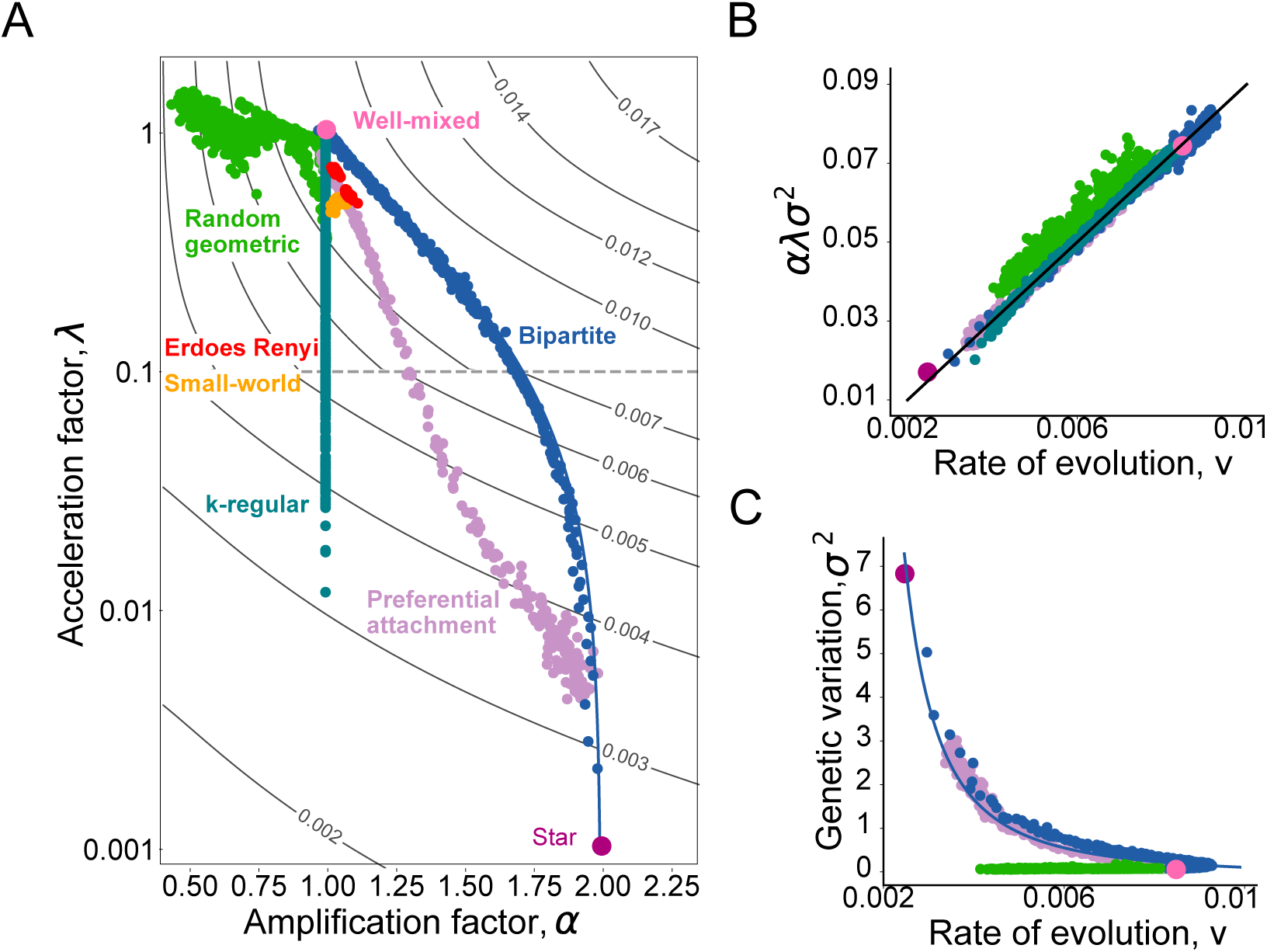
The rate of evolution and genetic variation across classes of population structures. Panel. **A.** The rate of evolution, as a function of the amplification and acceleration factor for graphs of size 1000. The mutation rate is set to 10*^−^*^4^ and *Ns* = 10. The black lines are contour lines of the rate of evolution using equation (9), and the dots represent individual networks of the families depicted. Network amplification factors empirically determined using equation (12). Network acceleration factors empirically determined using equation (13). **Panel B.** The relationship between rate of evolution and scaled mutational variation. The dots represent ensemble averages across at least 100 replicate Monte Carlo simulations per parameter set and network. The black line represents equation (8). Same parameters as **Panel A**. **Panel C.** The relationship between rate of evolution and mutational variation. Same parameters as **Panel A**.

We next investigate how the population structure reshapes the maintenance of genetic variation. According to Fisher’s fundamental theorem of natural selection, if the mutation rate is negligible, the relationship between speed of evolution and the variance in mutation number is given by *sV* = *s*^2^*σ*^2^ (Fisher, 1999). For network-structured populations, equation (8) gives us the new relationship between *V* and *σ*, which now also depends on the product *αλ* (Figure 7B). Controlling for this extra factor, in Figure 7C we observe that, for some structures, such as preferential attachment graphs and bipartite graphs, there is an inverse relationship between the rate of evolution and the amount of mutational variance. The majority of graphs in these families follow what is commonly accepted on the role of spatial structure: structure reduces the rate of evolution and increases genetic variation. This is because, for these graphs, the contribution from the acceleration factor outweighs the contribution from the amplification factor. Other structures, like the random geometric graphs, on the other hand, appear to only influence the rate of evolution and not significantly affect mutational variance (Figure 7C, the random geometric graphs in color green). However, upon closer look, for some of these geometric graphs, *σ*^2^ can be as low as 0.057, compared to *σ*^2^ = 0.0746 for well-mixed populations. This corresponds to a significant reduction in mutational variance (24%).

#### Rates of evolution and mutational variation across systems of organization

We next apply our model to study the rate of evolution in two natural systems with drastically different spatial architectures and organization, a cellular and a social system.

We first study the stem cell architectures of the bone marrow to understand spatial factors shaping the accumulation of somatic mutations and rates of neoplasm initiation. We use previously published datasets and inferred cellular networks, in total 16 cellular populations (Gomariz et al., 2018; Kuo et al., 2024; Kuo and Carja, 2024a). Every Hematopoietic Stem Cell (HSC) niche constitutes a node in the graph and an edge is added between two nodes if the distance between them is less than a cut-off radius, similar to the generation of a random geometric graph (Figure 8A). In Kuo et al. (2024) we have previously showed that, across a wide variety of parameters and regardless of the birth death process used, these networks are strong suppressors of selection, in the single mutation regime. Estimates of the mutation rate in this system, for mutations with selective advantage of 0.04 or higher, are around 4 × 10*^−^*^6^ per year, and HSCs are estimated to divide symmetrically every 4 years (upper-bound division time) (Watson et al., 2020).

**Figure 8:**
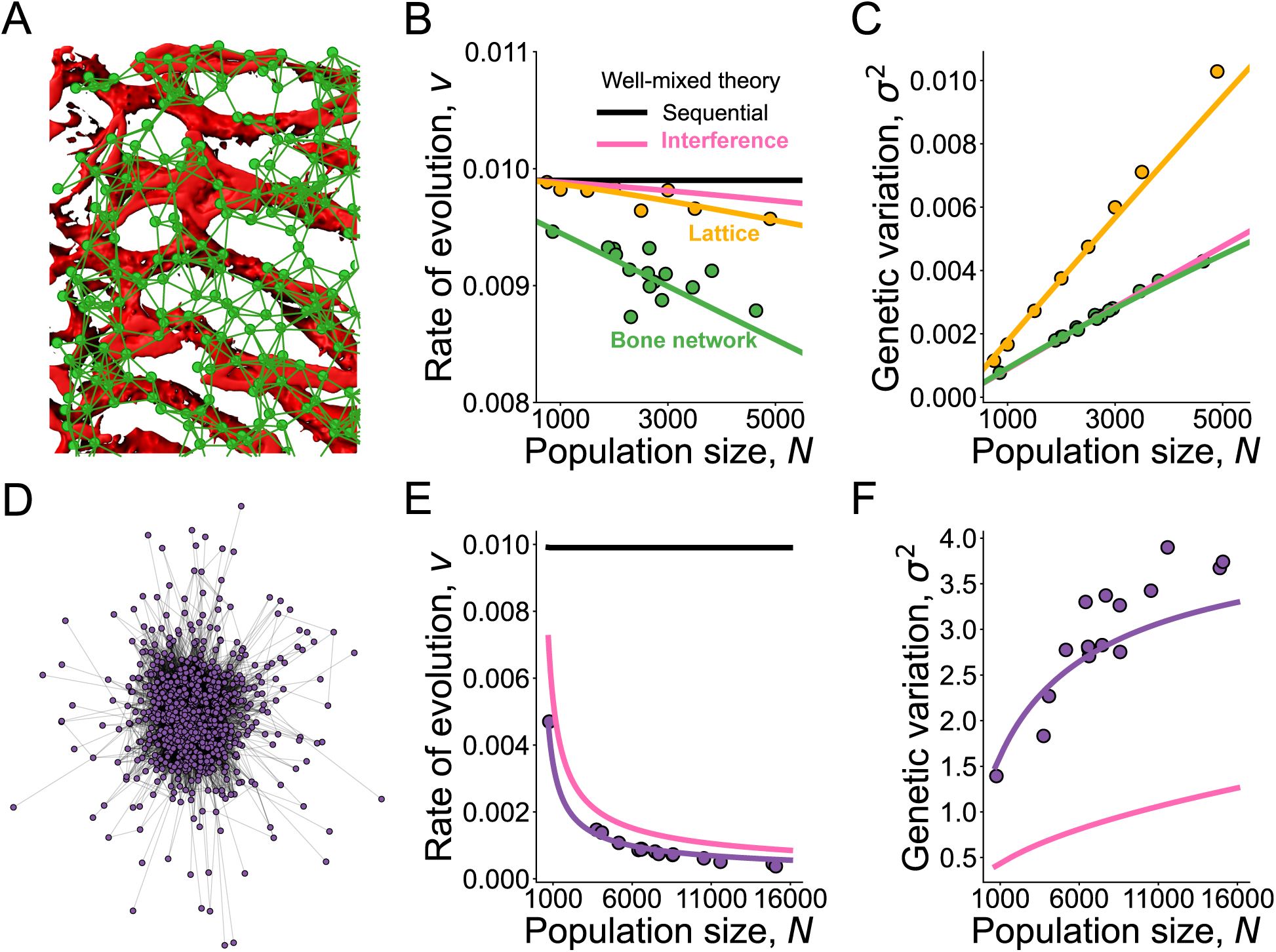
Model applications. Trade-offs between rate of evolution and genetic variation in cellular and social systems. Panel. **A.** Illustration of the hematopoietic stem cell niche architecture (for a full description see (Kuo et al., 2024)). **Panel B.** Rate of evolution in cellular networks of various sizes, *N*, between 1000 and 5000 cells. Here, *s* = 0.01, *U* = 10*^−^*^6^. Dots are simulation results using at least 1000 replicate Monte Carlo simulations per parameter set and network and lines are analytic approximations using equation (9). **Panel C.** Genetic variation in cellular networks. Same parameters as **Panel B**. **Panel D.** Illustration of a social network (here a Facebook network from Caltech, using published datasets from Rossi and Ahmed (2015)). **Panel E.** Rate of evolution in social networks of various sizes *N*. Here, *s* = 0.01, *U* = 10*^−^*^3^. Dots are simulation results using at least 1000 replicate Monte Carlo simulations per parameter set and network and lines are analytic approximations using equation (9). Purple line represents the population structure, while the pink line represents the well-mixed model as comparison. **Panel F.** Genetic variation in social networks. Same parameters as **E**. Social networks can significantly slow down the rate of cultural evolution, but also increase the amount of mutational variation, compared to well-mixed populations.

Using these parameters, we find that the bone marrow suppresses the rate of mutation accumulation, with the existence of prevalent clonal interference, indicated by the reduced rate of evolution compared to the estimate for an equivalent sequential fixation model (Figure 8B). While a lattice-based model also showed reduced rates of mutation accumulation, the inferred cellular network architectures decreased the rate of evolution by an even larger amount. In addition, they show very different behavior when it comes to the mutational variation in the population (Figure 8C). The bone marrow networks tend to have the same or lower mutation variation as the well-mixed population, in contrast to lattice-based populations, where the population topology promotes higher mutation variation. These results showcase that spatial heterogeneity in tissue architectures can reshape the rates of mutation accumulation and mutation variation in ways that can be overlooked by previous spatial models.

The second type of structures we consider are social networks and use previously published data on Facebook friendship networks, across 16 universities (Rossi and Ahmed, 2015). Since most of the social networks in the dataset are disconnected, we extract and use the largest connected component from each network (Figure 8D). We consider a simplified version of the model proposed by Kolodny et al. (2015), where the genome of an individual represents a repertoire of innovations or tools and study rates of cultural innovation accumulating in the population. We assume that new innovations enter the population independently of the current state of cultural diversity (like genetic mutations), with probability *U*. Kolodny et al. (2015) used *U* between 0.001 and 0.002. Here, we picked *U* = 0.001 and the Birth-death process on networks to represent the spread of the repertoire of tools. One would expect the topological structure of human interaction to maximize the rate of cultural evolution. However, Figure 8E shows the opposite—social networks slow down the rate of cultural evolution, but, interestingly, significantly increase the amount of mutational variation, compared to a well-mixed model (Figure 8F).

## Discussion

Here we use the mathematical representation of networks to study clonal interference and rates of evolution in populations with complex population structure. At a big picture level, prior modeling work done under assumptions of more regular and symmetric population architectures suggests that: 1) spatial structure increases the probability that established clones interfere with one another, 2) interference between established clones causes spatial populations to evolve slower than well-mixed populations, and 3) the relationship between the rate of evolution and genetic variance follows Fisher’s fundamental theorem of natural selection. We show that increasingly introducing more heterogeneity in the population structure considered can reshape these prior conclusions and lead to a much wider range of observed evolutionary outcome. Using simulations and analytic approximations we discuss how there exist two main evolutionary properties of spatial structure that drive the observed outcomes. We study their independent roles, as well as their interplay, in shaping evolutionary dynamics. Importantly, we find that, depending on their evolutionary properties, population structures can also lead to decreased rates of clonal interference, higher rates of mutation utilization and increased rates of evolution. For example, spatially heterogeneous graphs, such as those generated by sampling node positions from inhomogeneous distributions, are suppressors (networks with low amplification factor) and can reduce the probability that established clones interfere with one another.

The amount of clonal interference in the population can be quantified by mutation utilization, which measures how many established clones it takes to increase the average mutational count of the population by one and we show that suppressor topologies have a higher mutation utilization than well-mixed populations. Additionally, while previous theory suggests that more interference slows down evolution (Kryazhimskiy et al., 2012), here we discuss how there exist population structures with the right amount of heterogeneity that can strike a balance between the benefit of amplification and the costs of increased interference, achieving a higher rate of evolution than well-mixed populations in the interference regime. These amplifier topologies have a higher rate of evolution, even when there is more interference and we show that this happens when the benefit of having more mutants establish in the population outweighs the cost of interference.

In addition to looking at the mean number of mutations in the population, we also explore the distribution of the mutation count in the population using analytics and simulations. We use the variance of the distribution as measure of the width of the mutation count distribution. In well-mixed populations, the relationship between the rate of evolution and the variance in fitness obeys Fisher’s fundamental theorem of natural selection, *V* = *sσ*^2^ (Fisher, 1999). Otwinowski and Krug (2014) previously showed that Fisher’s fundamental theorem is no longer true on 1-dimensional lattices. Here, we derive the modified Fisher’s fundamental theorem, *V* = *αλsσ*^2^ for any population structure.

While our results in the limit of homogeneous spatial structure qualitatively agree with previous theory, it is worth discussing the slight quantitative differences. Tsimring et al. (1996) and Cohen et al. (2005) showed that the speed of evolution scales with *U* ^2^*^/^*^3^ in well-mixed populations, when mutation rate is high. Additionally, Martens et al. (2011) showed that the speed of evolution scales with *U* ^1^*^/^*^2^ for 1-dimensional lattices and *U* ^1^*^/^*^3^ for 2-dimensional lattices. This suggests that spatial structure affects the scaling relationship between speed of evolution and mutation rate. Our analytic results show that the scaling relationship for graph structured populations is the same as the well-mixed structure, with *V* ∼ *U* ^2^*^/^*^3^: the network structure only changes the scaling pre-factors. This difference in scaling comes from the diameter of the graph, i.e. the maximum shortest-path distance between two nodes of the graph. The *U* ^1^*^/^*^2^ and *U* ^1^*^/^*^3^ scaling in Martens et al. (2011) appears when the length of the lattice is large, larger than a characteristic length. We did not observe the same difference in scaling since many of the networks both in our simulations and in real-world systems have the small-world property, where the distance between any pair of nodes is smaller (Watts and Strogatz, 1998).

Our approach of distilling the role of population structure into the two evolutionary parameters of ‘space’, *α* and *λ*, retains sufficient information about the studied topology, is intuitively simple to understand, and produces very accurate predictions of the dynamics of mutation accumulation in real populations. To demonstrate this, we study the rate of evolution in two natural systems, a cellular and a social system. We first use previously studied datasets of cellular networks (Gomariz et al., 2018; Kuo et al., 2024) and study rates of mutation accumulation and leukemia initiation for the structured populations of stem cells in the bone marrow. Our results show that these cellular spatial architectures reduce the rate of neoplasm initiation under biologically relevant mutation rates, compared to well-mixed populations. In addition, these cellular networks do not increase mutational variation, unlike lattice-based models. These results highlight that spatial heterogeneity in tissue architectures can reduce the rates of mutation accumulation and mutation variation in ways that can be overlooked by previous spatial models.

We also use publicly available datasets of Facebook social connection networks (Rossi and Ahmed, 2015) and study how social structure shapes cultural accumulation of innovations in a population. In contrast to cellular structures, we find that social networks can promote maintenance of higher levels of cultural diversity, even though they can, similarly, reduce the rate of innovation accumulation. If we hypothesize that these networks are themselves shaped by natural selection, our results hint at a potential tug-of-war between optimizing for speed or variation in these systems. An aspect that is unique to culturally evolving systems, as opposed to somatic evolution, is the notion that truly groundbreaking innovations occur when individuals combine multiple low-level innovations (Basalla, 1988). Intuitively, successful recombination of existing innovations requires the maintenance of variability of innovations, and one can hypothesize that social structure might have had critical effects for the rise of cumulative culture. Derex and Boyd (2015) suggest that the rate of cultural accumulation may be highest in sizable and partially connected populations. This is also observed in our results in both artificial and empirical networks.

In the quest to balance model realism and analytical tractability, it is important to know when it is sufficient to investigate the role of population structure in the symmetric limit (where, on average, it is easier to obtain closed-form solutions), versus when it might be essential to consider the potentially more heterogenous and ‘messy’ structure of the system. This is requisite for forming sensible null expectations about experimental, observational, and genomic data. The models we develop here have their own limitations and strong assumptions, but they provide an initial unifying step towards understanding the role of population topology in shaping the dynamics of adaptation.

## Materials and methods

### List of network families used in the study

In this study, we explore well-known network families using built-in generators from NetworkX (Hagberg et al., 2008), as well as methods from Kuo et al. (2024) and Kuo and Carja (2024b) to design graphs that allow us to tune topological properties independently of each other, as detailed below.

### *k*-regular graphs

A *k*-regular graph is a graph where each node has the same number of neighbors, *k*. We generate *k*-regular graphs using built-in generators from NetworkX (Hagberg et al., 2008), with *k* ∈ {3, 4, 6, 8, 10}. For some of these graphs, to generate Figures 2, 3 and 4 we further tune the number of triangles and change their acceleration factor (while keeping mean degree constant at *k*), using methods perviously outlined in Kuo and Carja (2024b).

### Erdős Rényi random networks

We use the *G*(*n, M*) variant of Erdős Rényi model. The Erdős Rényi *G*(*n, M*) model uniformly samples a graph from the collection of all graphs which have n nodes and *M* edges. We generate Erdős Rényi random networks using built-in generators from NetworkX (Hagberg et al., 2008). Here, we sample 50 graphs with *M* = 3000 and 50 graphs with *M* = 4000.

### Preferential attachment graphs

For graphs with preferential attachment, nodes are added sequentially starting from one initial node until the population reaches size *N*. Each new node is added to the network and connected to other individuals with a probability proportional to the individual’s current degree to the power of a given parameter *β*. Using *k* and *β,* this family of graphs allows for straightforward independent tuning of mean and variance in degree. We generate these networks using NetworkX (Hagberg et al., 2008) with *k* = 4, and *β* ∈ [−5, 5].

### Small world networks

We generate small world networks using built-in generators from NetworkX (Hagberg et al., 2008) and the Watts–Strogatz model with parameters: number of nodes *N*, the mean degree *k*, and rewire probability *p*. We use *N* = 1000*, k* = 4, and *p* ∈ [0, 1].

### Bipartite graphs

Bipartite graphs have two node sets *n*_1_ and *n*_2_. Edges in the graph only connect nodes from opposite sets. We generate bipartite graphs using built-in generators from NetworkX (Hagberg et al., 2008). We vary *n*_1_ from 1 to 500, and *n*_2_ = 1000 − *n*_1_.

### Random geometric graphs

The random geometric graph model places *N* nodes at random following a probability distribution. Two nodes are joined by an edge if the distance between the nodes is below a predefined cut-off radius. For the graphs in Figure 5, we place nodes in 2-dimensional space. We draw the spatial location from two distributions. We first use the Gaussian distribution, where for the *x*-position, *n*_1_ = 500, 600, 700, 800, 900 nodes are drawn from a normal distribution N (0, 1), and *n*_2_ = 1000 − *n*_1_ nodes are drawn from N (0.5, 1). The *y*-position is drawn from N (0, 4). We also use the uniform distribution, where for the *x*-position, *n*_1_ = 500, 600, 700, 800, 900 nodes are drawn from a normal distribution U(0, 1), and *N*_2_ = 1000 − *n*_1_ nodes are drawn from N (0.5, 1.5). The *y*-position is drawn from U(0, 1). We used cut-off radius ranging from 0 to 4.

### Star graphs

The star graph consists of one center node connected to *N* − 1 outer nodes. We generate star graphs using built-in generators from NetworkX.

### Computing the amplification and acceleration parameters for a given network

Here we show that the amplification and acceleration factors of a network, computed from single mutation analyses, if used together, can be predictive of the landscape crossing behavior of network structured populations. In previous work, we showed how to analytically compute the amplification and acceleration factors (Kuo et al., 2024; Kuo and Carja, 2024b). The approximation for the amplification factor from Kuo et al. (2024) is very accurate for amplifiers and can slightly deviate from the true amplification factor for suppressors. Similarly, the approximations described in Kuo and Carja (2024b) for computing the acceleration factor are very accurate if triangles in the network are distributed equally amongst the triplet types.

These factors can also be computed empirically for all network families and types using single-mutation simulations, as detailed below. Using the Birth-death process, and starting with one single mutant with fitness (1 + *s*) invading a population of wild-type individuals with fitness 1, the amplification factor can be computed using the definition from (Lieberman et al., 2005) and solving

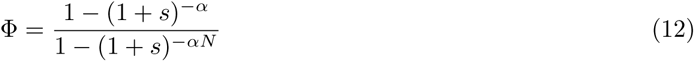

for *α*, where Φ is the mutant’s fixation probability.

For graphs in this paper, we estimate *α* using *s* = 0.01. When graph size is *N* = 1000, *Ns* = 10. This value is large enough such that the difference in fixation probability compared to the well-mixed model is big enough, while also small enough to be in the constant regime.

For the acceleration factor, we use the following definition

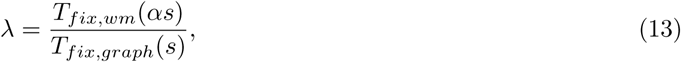

which is the ratio of conditional mean fixation time for an equivalent mutant on the well-mixed population (with selection coefficient *αs*) and the one on the network population with mutant selection coefficient *s* (Kuo and Carja, 2024b). For the graphs presented here, we evaluate *λ* at *s* = 0, since there is a closed-form expression for *T_fix,wm_*(*s* = 0), (*N* − 1) generations (Ewens, 2004). For weak *s, λ* varies no more than O(*s*).

### Data and code availability

Custom scripts were used for the simulations and data analyses. All C++ and Python simulation code, as well as the code to generate the graphs used in this study is available on Github at XXX (repository to be made public upon acceptance and publication of the manuscript). All packages used for analysis and visualization are open-source.

## Acknowledgments

This research was done using resources provided by the Open Science Grid, which is supported by the National Science Foundation award 1148698, and the U.S. Department of Energy’s Office of Science.

## Funding

We gratefully acknowledge support from the NIH National Institute of General Medical Sciences (award no. R35GM147445) and from the NIH T32 training grant (no. T32 EB009403).

## Conflicts of interest

The authors declare no conflicts of interest.

## Notes

### Competing Interest Statement

The authors have declared no competing interest.

## References

Ben Adlam, Krishnendu Chatterjee, and Martin A Nowak. Amplifiers of selection. Proceedings of the Royal Society A: Mathematical, Physical and Engineering Sciences, 471(2181):20150114, 2015.

Benjamin Allen, Christine Sample, Patricia Steinhagen, Julia Shapiro, Matthew King, Timothy Hedspeth, and Megan Goncalves. Fixation probabilities in graph-structured populations under weak selection. PLoS Computational Biology, 17(2):e1008695, 2021.

Tibor Antal, Sidney Redner, and Vishal Sood. Evolutionary dynamics on degree-heterogeneous graphs. Physical Review Letters, 96(18):188104, 2006.

Armen S Asratian, Tristan MJ Denley, and Roland Häggkvist. Bipartite graphs and their applications, volume 131. Cambridge University Press, 1998.

Albert-László Barabási and Réka Albert. Emergence of scaling in random networks. Science, 286(5439): 509–512, 1999.

George Basalla. The evolution of technology. Cambridge University Press, 1988.

Francis Blokzijl, Joep De Ligt, Myrthe Jager, Valentina Sasselli, Sophie Roerink, Nobuo Sasaki, Meritxell Huch, Sander Boymans, Ewart Kuijk, Pjotr Prins, et al. Tissue-specific mutation accumulation in human adult stem cells during life. Nature, 538(7624):260–264, 2016.

Timothée Bonnet, Michael B Morrissey, Pierre De Villemereuil, Susan C Alberts, Peter Arcese, Liam D Bailey, Stan Boutin, Patricia Brekke, Lauren JN Brent, Glauco Camenisch, et al. Genetic variance in fitness indicates rapid contemporary adaptive evolution in wild animals. Science, 376(6596):1012–1016, 2022.

Éric Brunet, Igor M Rouzine, and Claus O Wilke. The stochastic edge in adaptive evolution. Genetics, 179 (1):603–620, 2008.

Oana Carja, Uri Liberman, and Marcus W Feldman. Evolution in changing environments: Modifiers of mutation, recombination, and migration. Proceedings of the National Academy of Sciences, 111(50):17935– 17940, 2014.

Joshua L Cherry and John Wakeley. A diffusion approximation for selection and drift in a subdivided population. Genetics, 163(1):421–428, 2003.

Elisheva Cohen, David A Kessler, and Herbert Levine. Front propagation up a reaction rate gradient. Physical Review E, 72(6):066126, 2005.

James F Crow, Motoo Kimura, et al. An introduction to population genetics theory. An introduction to population genetics theory, 1970.

Maxime Derex and Robert Boyd. The foundations of the human cultural niche. Nature Communications, 6 (1):8398, 2015.

Michael M Desai and Daniel S Fisher. Beneficial mutation–selection balance and the effect of linkage on positive selection. Genetics, 176(3):1759–1798, 2007.

Paul Erdős and Alfréd Rényi. On the evolution of random graphs. Publ. Math. Inst. Hung. Acad. Sci, 5(1): 17–60, 1960.

Warren John Ewens. Mathematical population genetics: theoretical introduction, volume 27. Springer, 2004.

R.A. Fisher. The genetical theory of natural selection. Oxford University Press, 1930.

Ronald Aylmer Fisher. The genetical theory of natural selection: a complete variorum edition. Oxford University Press, 1999.

Marcus Frean, Paul B Rainey, and Arne Traulsen. The effect of population structure on the rate of evolution. Proceedings of the Royal Society B: Biological Sciences, 280(1762):20130211, 2013.

Crispin Gardiner. Stochastic methods, volume 4. Springer Berlin, 2009.

Philip J Gerrish and Richard E Lenski. The fate of competing beneficial mutations in an asexual population. Genetica, 102:127–144, 1998.

Alvaro Gomariz, Patrick M Helbling, Stephan Isringhausen, Ute Suessbier, Anton Becker, Andreas Boss, Takashi Nagasawa, Grégory Paul, Orcun Goksel, Gábor Székely, et al. Quantitative spatial analysis of haematopoiesis-regulating stromal cells in the bone marrow microenvironment by 3D microscopy. Nature Communications, 9(1):2532, 2018.

Isabel Gordo and Paulo RA Campos. Adaptive evolution in a spatially structured asexual population. Genetica, 127:217–229, 2006.

Aric Hagberg, Pieter Swart, and Daniel S Chult. Exploring network structure, dynamics, and function using NetworkX. Technical report, Los Alamos National Lab.(LANL), Los Alamos, NM (United States), 2008.

Oskar Hallatschek. The noisy edge of traveling waves. Proceedings of the National Academy of Sciences, 108 (5):1783–1787, 2011.

Laura Hindersin and Arne Traulsen. Counterintuitive properties of the fixation time in network-structured populations. Journal of The Royal Society Interface, 11(99):20140606, 2014.

Laura Hindersin and Arne Traulsen. Most undirected random graphs are amplifiers of selection for birth-death dynamics, but suppressors of selection for death-birth dynamics. PLoS Computational Biology, 11 (11):e1004437, 2015.

Laura Hindersin, Marius Möller, Arne Traulsen, and Benedikt Bauer. Exact numerical calculation of fixation probability and time on graphs. Biosystems, 150:87–91, 2016.

Motoo Kimura. The number of heterozygous nucleotide sites maintained in a finite population due to steady flux of mutations. Genetics, 61(4):893, 1969.

Motoo Kimura and George H Weiss. The stepping stone model of population structure and the decrease of genetic correlation with distance. Genetics, 49(4):561, 1964.

Oren Kolodny, Nicole Creanza, and Marcus W Feldman. Evolution in leaps: the punctuated accumulation and loss of cultural innovations. Proceedings of the National Academy of Sciences, 112(49):E6762–E6769, 2015.

Sergey Kryazhimskiy, Daniel P Rice, and Michael M Desai. Population subdivision and adaptation in asexual populations of Saccharomyces cerevisiae. Evolution, 66(6):1931–1941, 2012.

Yang Ping Kuo and Oana Carja. Evolutionary graph theory beyond single mutation dynamics: on how network structured populations cross fitness landscapes. Genetics, page iyae055, 2024a.

Yang Ping Kuo and Oana Carja. Evolutionary graph theory beyond pairwise interactions: Higher-order network motifs shape times to fixation in structured populations. PLOS Computational Biology, 20(3): e1011905, 2024b.

Yang Ping Kuo, César Nombela-Arrieta, and Oana Carja. A theory of evolutionary dynamics on any complex population structure reveals stem cell niche architecture as a spatial suppressor of selection. Nature Communications, 15(1):4666, 2024.

Erez Lieberman, Christoph Hauert, and Martin A Nowak. Evolutionary dynamics on graphs. Nature, 433 (7023):312–316, 2005.

Peter V Markov, Mahan Ghafari, Martin Beer, Katrina Lythgoe, Peter Simmonds, Nikolaos I Stilianakis, and Aris Katzourakis. The evolution of SARS-CoV-2. Nature Reviews Microbiology, 21(6):361–379, 2023.

Erik A Martens and Oskar Hallatschek. Interfering waves of adaptation promote spatial mixing. Genetics, 189(3):1045–1060, 2011.

Erik A Martens, Rumen Kostadinov, Carlo C Maley, and Oskar Hallatschek. Spatial structure increases the waiting time for cancer. New Journal of Physics, 13(11):115014, 2011.

Takeo Maruyama. Effective number of alleles in a subdivided population. Theoretical Population Biology, 1 (3):273–306, 1970.

Travis Monk, Peter Green, and Mike Paulin. Martingales and fixation probabilities of evolutionary graphs. *Proceedings of the Royal Society A: Mathematical*, Physical and Engineering Sciences, 470(2165):20130730, 2014.

H. Muller. Some genetic aspects of sex. Am. Nat., 66:2118–138, 1932.

Jakub Otwinowski and Joachim Krug. Clonal interference and muller’s ratchet in spatial habitats. Physical Biology, 11(5):056003, 2014.

Andreas Pavlogiannis, Josef Tkadlec, Krishnendu Chatterjee, and Martin A Nowak. Amplification on undirected population structures: comets beat stars. Scientific Reports, 7(1):82, 2017.

Mathew Penrose et al. Random geometric graphs, volume 5. Oxford University Press, 2003.

Gladys Y. P. Poon, Caroline J. Watson, Daniel S. Fisher, and Jamie R. Blundell. Synonymous mutations reveal genome-wide levels of positive selection in healthy tissues. Nature Genetics, 53(11):1597–1605, 2021.

Ryan A. Rossi and Nesreen K. Ahmed. The network data repository with interactive graph analytics and visualization. In AAAI, 2015. URL https://networkrepository.com.

Igor M Rouzine, John Wakeley, and John M Coffin. The solitary wave of asexual evolution. Proceedings of the National Academy of Sciences, 100(2):587–592, 2003.

Igor M Rouzine, Éric Brunet, and Claus O Wilke. The traveling-wave approach to asexual evolution: Muller ratchet and speed of adaptation. Theoretical Population Biology, 73(1):24–46, 2008.

David M Schneider, Ayana B Martins, and Marcus AM de Aguiar. The mutation–drift balance in spatially structured populations. Journal of Theoretical Biology, 402:9–17, 2016.

Nikhil Sharma and Arne Traulsen. Suppressors of fixation can increase average fitness beyond amplifiers of selection. Proceedings of the National Academy of Sciences, 119(37):e2205424119, 2022.

Montgomery Slatkin. Fixation probabilities and fixation times in a subdivided population. Evolution, pages 477–488, 1981.

Jakub Svoboda, Soham Joshi, Josef Tkadlec, and Krishnendu Chatterjee. Amplifiers of selection for the Moran process with both birth-death and death-birth updating. PLOS Computational Biology, 20(3): e1012008, 2024.

Josef Tkadlec, Andreas Pavlogiannis, Krishnendu Chatterjee, and Martin A Nowak. Population structure determines the tradeoff between fixation probability and fixation time. Communications Biology, 2(1): 1–8, 2019.

Josef Tkadlec, Andreas Pavlogiannis, Krishnendu Chatterjee, and Martin A Nowak. Limits on amplifiers of natural selection under death-birth updating. PLoS Computational Biology, 16(1):e1007494, 2020.

Lev S Tsimring, Herbert Levine, and David A Kessler. RNA virus evolution via a fitness-space model. Physical Review Letters, 76(23):4440, 1996.

John Wakeley. Segregating sites in Wright’s island model. Theoretical Population Biology, 53(2):166–174, 1998.

Caroline J Watson, AL Papula, Gladys YP Poon, Wing H Wong, Andrew L Young, Todd E Druley, Daniel S Fisher, and Jamie R Blundell. The evolutionary dynamics and fitness landscape of clonal hematopoiesis. Science, 367(6485):1449–1454, 2020.

Duncan J Watts and Steven H Strogatz. Collective dynamics of small world networks. Nature, 393(6684): 440–442, 1998.

Bernard M Waxman. Routing of multipoint connections. IEEE journal on Selected Areas in Communications, 6(9):1617–1622, 1988.

Michael C Whitlock. Fixation probability and time in subdivided populations. Genetics, 164(2):767–779, 2003.

Michael C Whitlock and NH Barton. The effective size of a subdivided population. Genetics, 146:427–441, 1997.

Sewall Wright. Isolation by distance. Genetics, 28(2):114, 1943.

