## Supplementary Information for "Clonal interference, genetic variation and the speed of evolution in structured populations"

**4 Supplementary Figures** **25**

### 1 Deriving the equations governing the evolutionary dynamics

In this section, we describe our analytical approach to the problem of mutation accumulation for graph structured populations. We first show how we derive the Kolmogorov backward equation describing the dynamics of mutation accumulation and then discuss how to derive the approximations used in the main text, for the two quantities of interest: the rate of evolution, and the genetic variance in mutation number.

At a big picture level, to make analytical headway, we will use the diffusion approximation. Direct application of the diffusion approximation, however, is difficult, due to the complex dynamics that can occur if the population structure is a graph. In a well-mixed population (or on the complete graph), keeping track of the different mutant frequencies in the population is sufficient for determining and understanding the evolutionary fate of the population. However, for a population with heterogeneous spatial structure, what makes the problem particularly challenging is that keeping track of the population-level mutational frequencies is no longer sufficient, as the replacement between different mutants depends on the frequencies of different types of nodes (and their degrees), as well as the frequencies of the different types of edges (and what types of nodes they connect) in the network population.

The approach we take, and the core idea behind our approximation, is that we can greatly reduce the complexity of the problem by using the node degree distribution of the network, and only keeping track of mutant frequencies for all groups of nodes of the same degree (instead of the mutant state of all possible nodes, which would make the analytics intractable for large networks). While the degree distribution might not uniquely represent the network and some of the graph information is lost, this approach nonetheless greatly reduces the number of possible states in the Moran model. **Another way to look at this: by**

**grouping together nodes of the same degree and tracking mutant frequency changes on these groups of nodes, we essentially transform the problem into a finite island population type model.**

To start with, we first show that if we assume nodes of the same degree in the network to be 'evolutionarily' identical, we can derive the expected change in the $l$-mutant lineage frequency for each group of nodes of equal degree. To this end, we first observe that, since replacements between two lineages, $l$-mutants and $m$-mutants, depend on the number of edges between these two mutant types, we can write the expected change in the $l$-mutant frequencies at nodes of the same degree as a function of the frequencies of the different edge types possible in the graph.

By then deriving the expected change in edge type frequencies, if we assume weak selection and weak mutation, we find that the mutant frequencies for each group of nodes with distinct degrees is the same for each l-mutant. We define this quasi-equilibrium node frequency as $x_l$, which depends on the initial mutant placements on the network. The quasi-equilibrium edge frequencies also depend on $x_l$. Selection and mutation, however, will slowly alter this quasi-equilibrium. (Note that we first discussed this quasi-equilibrium and used this approach in [Kuo and Carja, 2024a], a simpler two-mutant problem.)

In **Supplementary Material Sections 1.3-1.6**, we then write the expected change in $x_l$ due to selection and mutation. We then write the variance and covariance in the change of $x_l$ and finally, in the limit of weak selection $s \ll 1$, weak mutation $U \ll 1$, and large population size $1/N \ll 1$, we write the Kolmogorov backward equation for the network population dynamics. At a big picture level, we show that this equation simplifies to the equation in the well-mixed population limit, with the appropriate parameter stretches given by the two evolutionary network properties, amplification and acceleration.

#### 1.1 Deconstructing the dynamics into its components: network structure, selection, mutation and drift

To study how heterogenous population structure shapes patterns of genetic variation and long-term rates of adaptation in the concurrent mutations regime, we study an asexual population of $N$ haploid individuals. We use the infinite sites mutation model [Kimura, 1969], and assume that:

1. there are $b$ beneficial sites where mutations can occur,
2. $b$ is assumed to be very large such that no two mutations occur at the same site, and
3. there is no recombination.

New beneficial mutations appear at a rate $\mu$ per site and we study a model in which each beneficial mutation has the same effect, $s$, on fitness (i.e., each step uphill is of the same size). The total beneficial mutation rate per replication is therefore $U = \mu b$. There are no epistatic interactions between mutations, such that fitness is multiplicative and an individual with $l$ mutations has reproductive fitness of $(1+s)^l \approx 1 + ls$, for $s \ll 1$. Let us denote by $n_l$ and $p_l$ as the number and frequency of $l$-mutant individuals in the population.

Spatial structure is represented by a graph. We denote by $n_i$ and $p_i$ the number and frequency of nodes of degree $i$, and by $n_{ij}$ and $p_{ij}$ the number and frequency of edges that connect nodes of degree $i$ to nodes of degree $j$. Let $n_{i,l}$ denote the number of l-mutants that occupy nodes of degree $i$ and $n_{ij,lm}$ denote the number of edges that connect nodes $l_i$ and $m_j$, i.e. nodes of degree $i$ containing the $l$-mutant with nodes of degree $j$ containing the $m$-mutant. Similarly, we define $n_{ij,ll}$ and $n_{ij,mm}$. We also define frequencies $p_{i,l}$ as $n_{i,l}$ divided by the total population size and $p_{ij,lm}$ as $n_{ij,lm}$ divided by the total number of edges $\sum n_{ij}$.

We observe that $l_i$ will decrease by one whenever a $m$-mutant reproduces and replace a $l$-mutant on node with degree $i$. The probability that a $m$-mutant occupying a node of degree $j$ replaces a $l$-mutant occupying a node of degree $i$ is the probability of selecting a node $m_j$ to reproduce and a neighboring node $l_i$ to die. This probability is given by

$$\begin{aligned} P(m_j \to l_i) &= \frac{(1+s)^m}{Nw} n_{j,m} \frac{n_{ij,lm}}{j n_{j,m}} \\ &= \frac{(1+s)^m}{Nwj} n_{ij,lm}. \end{aligned} \tag{1}$$

Here, $w = \sum_l (1+s)^l p_l$ represents the mean fitness of the population.

Using this probability, we calculate the expected change in the numbers of different node types, over one update, as

$$m_j \to l_i : \quad \Delta n_{i,l} = -1 \quad \mathbb{E}[\Delta n_{i,l} \text{ due to } m_j \to l_i] = -\frac{(1+s)^m}{Nwj} n_{ij,lm} \tag{2}$$

$$l_j \to m_i : \quad \Delta n_{i,l} = +1 \quad \mathbb{E}[\Delta n_{i,l} \text{ due to } l_j \to m_i] = +\frac{(1+s)^l}{Nwj} n_{ij,ml}. \tag{3}$$

Summing the two expressions over all possible $m \neq l$, we compute the total expected change in the $l$-mutant numbers on nodes of fixed degree $i$, $n_{i,l}$, over one update and, for now, assuming no further mutation:

$$\mathbb{E}[\Delta n_{i,l}, \text{ no mutation}] = \sum_{j,m \neq l} \frac{1}{Nwj} \left[ (1+s)^l n_{ij,ml} - (1+s)^m n_{ij,lm} \right]. \tag{4}$$

Even with our approach of using the node degree distribution to simplify the problem, equation (4) shows that, in order to calculate the number of $l$-mutants on the graph, we need to consider all combinations of node degrees with different mutation numbers and all combinations of edge types with pairs of mutation numbers.

We rearrange equation (4) to simplify it. Under a weak selection assumption, $Ns \ll 1$, we expand all the terms containing $s$ and write

$$\begin{aligned}
\mathbb{E}[\Delta n_{i,l}, \text{ no mutation}] &= \sum_{j,m \neq l} (1 - \bar{s} + \mathcal{O}(s^2)) \frac{1}{Nj} \left[ (1+s)^l n_{ij,ml} - (1+s)^m n_{ij,lm} \right] \\
&= \sum_{j,m \neq l} \frac{1}{Nj} \left[ (1 + ls - \bar{s} + \mathcal{O}(s^2)) n_{ij,ml} - (1 + ms - \bar{s} + \mathcal{O}(s^2)) n_{ij,lm} \right] \\
&= \sum_{j} \frac{1}{Nj} (n_{ij,Ll} - n_{ij,lL}) + \sum_{j,m \neq l} \frac{1}{Nj} \left[ (ls - \bar{s}) n_{ij,ml} - (ms - \bar{s}) n_{ij,lm} \right] + \mathcal{O}(s^2) \\
&= \sum_{j} \frac{1}{Nj} (n_{ij,Ll} - n_{ij,lL}) + \frac{1}{N} \mathcal{O}(s).
\end{aligned} \tag{5}$$

Here, we expanded $w^{-1} = 1 - \bar{s} + \mathcal{O}(s^2)$, where $1 + \bar{s}$ is the mean population fitness. We use $L$ to represent all $m$-mutants with number of mutations $m \neq l$, such that $n_{ij,Ll} = \sum_{m \neq l} n_{ij,ml}$. This effectively helps clarify equation (4), by explicitly writing out terms that involve the $l-$mutant and terms without it.

These derivations above assumed we are in a no mutation limit. Let us now consider non-negligible mutation. The probability that there is a mutation from the $l$-mutant into the $(l+1)$-mutant is given by

$$P(\Delta n_{i,l} = -1, \text{ due to mutation}) = \frac{U}{N} n_{i,l}. \tag{6}$$

This change in the $l$-mutant numbers due to mutation is therefore given by

$$\mathbb{E}[\Delta n_{i,l}, \text{ due to mutation}] = \frac{U}{N} (n_{i,l-1} - n_{i,l}). \tag{7}$$

Summing these expectations, with and without the possibility of mutation, we can compute the total expected change in the $l$-mutant numbers on nodes of fixed degree $i$, $n_{i,l}$, over one update, as

$$\mathbb{E}[\Delta n_{i,l}] = \sum_j \frac{1}{Nj}\left(n_{ij,Ll} - n_{ij,lL}\right) + \frac{1}{N}\mathcal{O}(s) + \frac{U}{N}(n_{i,l-1} - n_{i,l}). \tag{8}$$

The change in the mutant numbers over one generation, as $N \to \infty$ and $\Delta T = 1/N \to 0$, can be written as

$$\frac{d}{dt}n_{i,l} = \underbrace{\sum_j \frac{1}{j}\left(n_{ij,Ll} - n_{ij,lL}\right) +}_{\text{Network structure}} \underbrace{\mathcal{O}(s)}_{\text{Network-dependent selection}} \underbrace{+\, U\left(n_{i,l-1} - n_{i,l}\right)}_{\text{Mutation}}. \tag{9}$$

The change in the $l$-mutant frequency over time therefore critically depends on the frequencies of the different edge types in the network and, to get a full description of the dynamics, we need to calculate this change in edge types over one update.

In what follows, we analyze the different terms in the above equation, sequentially.

#### 1.2 Analysis of the network structure term

As mentioned above, in a well-mixed population or on the complete graph, keeping track of the $l$-mutant frequencies on all nodes of degree $i$ (in effect, one single frequency for each distinct $l$ lineage) is sufficient for determining the evolutionary fate of the population. However, what makes the problem we solve particularly challenging is that here, the replacement between different mutants depends on the frequency of edges that connect the two in the network population, in other words depends on how many edges exist between $l$-mutants and $L$-mutants.

For simplicity and clarity, in what follows let us assume $s \ll 1$ and $U \ll 1$. The edge count $n_{ij,lL}$ can change in three types of events. The first two types of events are represented by a replacement event between a $l$-mutant occupying a node of degree $i$ and a $L$-mutant occupying a node of degree $j$, and vice versa. The expected update in $n_{ij,lL}$ is given by

$$\begin{aligned} l_i \to L_j: \quad \Delta n_{ij,lL} = -1 \quad \mathbb{E}[\Delta n_{ij,lL} \text{ due to } l_i \to L_j] &= -\frac{1}{Ni}n_{ij,lL} + \mathcal{O}(s) + \mathcal{O}(U) \\ L_j \to l_i: \quad \Delta n_{ij,lL} = -1 \quad \mathbb{E}[\Delta n_{ij,lL} \text{ due to } L_j \to l_i] &= -\frac{1}{Nj}n_{ij,lL} + \mathcal{O}(s) + \mathcal{O}(U). \end{aligned} \tag{10}$$

Here, we again use $L$ to represent all $m$-mutants with number of mutations $m \neq l$. One might be misled to think that the edge updates only depend on the total frequency $l$-mutants and $L$-mutants, and the actual distribution of $m$-mutants on the different nodes of the network does not matter. However, the dynamics of edge frequencies does depends on $m$ mutant values, since it affects the selective advantage each mutant class has. There is no discrepancy here because the effect of the individual $m$-mutants is contained in the $\mathcal{O}(s)$ term. For now, we are only explicitly writing out terms that are independent of $s$. Indeed, in the limit of $s \to 0$, the actual value of $m$ no longer matters, because all mutants would have fitness equal to one. For these other tems, we can effectively treat all $m \neq l$ as the same $L$-mutant.

The combined expected change under these two events is given by

$$\mathbb{E}[\Delta n_{ij,lL}, i \leftrightarrow j] = -\frac{1}{N}\left(\frac{1}{i} + \frac{1}{j}\right) n_{ij,lL} + \mathcal{O}(s) + \mathcal{O}(U). \tag{11}$$

The third event type is when a third individual occupying a neighbor node of degree $k$ replaces a node of degree $i$ or $j$. Under this scenario, there are four possible ways for the heterogeneous edge to change in number. For example, the probability of $l_k \to L_j l_i$ ( $l_k L_j l_i$ turning into $l_k l_j l_i$), is

$$P(l_k \to L_j l_i) = \frac{1}{Nk} n_{kj,lL} \frac{n_{kji,lLl}}{n_{kj,lL}(j-1)} + \mathcal{O}(s) + \mathcal{O}(U). \tag{12}$$

Here, the first part of the first term is the probability that $l_k$ is selected to replace $L_j$. The second fraction in the first term is the frequency of the $l_k L_j l_i$ triplet in all triplets containing the edge $l_k L_j$. This frequency is equal to the probability that an $l_k L_j$ edge forms an $l_k L_j l_i$ triplet, where $n_{kji,lLl}$ is the number of $l_k L_j l_i$ triplets. The probability in equation (12) is over one triplet, but there are $(j-1)$ total triplets centered around $L_j$. To make analytical headway, we use pair approximations and assume that the formation of $l_k L_j l_i$ triplets is independent to other triplets in the network [Pugliese and Castellano, 2009, Kuo and Carja, 2024b].

Under this independence assumption, the number of $l_k L_j l_i$ triplets is $(j-1)$ times the second fraction in the first term of equation (12). Therefore the expected transition from $l_k L_j l_i$ to $l_k l_j l_i$ is $(j-1)$ times equation (12), which is given by

$$\begin{aligned} l_k \to L_j l_i: &\quad \mathbb{E}[\Delta n_{ij,lL}, \text{ due to } l_k \to L_j l_i] = -n_{kji,lLl}/(Nk) + \mathcal{O}(s) + \mathcal{O}(U), \\ L_k \to l_j l_i: &\quad \mathbb{E}[\Delta n_{ij,lL}, \text{ due to } L_k \to l_j l_i] = +n_{kji,Lll}/(Nk) + \mathcal{O}(s) + \mathcal{O}(U). \end{aligned} \tag{13}$$

The expected change is then given by

$$\mathbb{E}[\Delta n_{ij,lL}, \text{ due to } k \to ji] = \sum_k \frac{1}{Nk}(n_{kji,Lll} - n_{kji,lLl}) + \mathcal{O}(s) + \mathcal{O}(U). \tag{14}$$

Similarly, we can write

$$\begin{aligned} l_k \to L_i L_j : \quad & \mathbb{E}[\Delta n_{ij,lL}, \text{ due to } l_k \to L_i L_j] = +n_{kij,lLL}/(Nk) + \mathcal{O}(s) + \mathcal{O}(U) \\ L_k \to l_i L_j : \quad & \mathbb{E}[\Delta n_{ij,lL}, \text{ due to } L_k \to l_i L_j] = -n_{kij,LlL}/(Nk) + \mathcal{O}(s) + \mathcal{O}(U). \end{aligned} \tag{15}$$

The expected change under these two events is given by

$$\mathbb{E}[\Delta n_{ij,lL} \text{ due to } k \to ij] = \sum_k \frac{1}{Nk}(n_{kij,lLL} - n_{kij,LlL}) + \mathcal{O}(s) + \mathcal{O}(U). \tag{16}$$

Summing (11), (14), and (16) allows us to write out the deterministic edge dynamics, assuming no drift for now. As $N \to \infty$ and $\Delta T = 1/N \to 0$,

$$\frac{d}{dt} n_{ij,lL} = -\left(\frac{1}{i} + \frac{1}{j}\right) n_{ij,lL} + \sum_k \frac{1}{k}(n_{kji,Lll} - n_{kji,lLl} + n_{kij,lLL} - n_{kij,LlL}) + \mathcal{O}(s) + \mathcal{O}(U). \tag{17}$$

We call equation (17), with $s = 0$ and $\mu = 0$, the neutral edge dynamics. If the selection and mutation terms are small, then the network term dominates. In this case, we can assume that the neutral edge dynamics has enough time to reach to an equilibrium before selection and mutation can have a non-negligible effect on the population.

This quasi-equilibrium point (in node and edge dynamics) is then calculated by setting the right-hand side of equations (9) and (17) to 0 and solving

$$\begin{cases} 0 = \sum_j \frac{1}{j}(n_{ij,Ll} - n_{ij,lL}), & \text{node equilibrium;} \\ 0 = -\left(\frac{1}{i} + \frac{1}{j}\right) n_{ij,lL} + \sum_k \frac{1}{k}(n_{kji,Lll} - n_{kji,lLl} + n_{kij,lLL} - n_{kij,LlL}), & \text{edge equilibrium.} \end{cases} \tag{18}$$

Using the framework and results from Kuo and Carja [2024b] (see supplementary equation (32)) and Kuo and Carja [2024a], we find that the equilibrium distribution of the $l$-mutant node is given by

$$x_l = \frac{1}{\mathbb{E}[i^{-1}]} \sum_i p_i p_{l|i} \frac{1}{i}, \quad \text{where} \quad \mathbb{E}[i^{-1}] = \left(\sum_i p_i \frac{1}{i}\right)^{-1}, \tag{19}$$

where $p_{l|i} = \frac{p_{i,l}}{p_i}$ is the probability of the mutant type being $l$ conditional on node being of degree $i$. We can write a similar equation for $p_{L|i}$. Additionally, the equilibrium distributions of edges linking mutants are given by

$$\begin{cases} x_{ij,LL} = p_{ij} p_{L|i} (1 - c_{ij} p_{l|j}), \\ x_{ij,Ll} = p_{ij} c_{ij} p_{L|i} p_{l|j}, \\ x_{ij,lL} = p_{ij} c_{ij} p_{l|i} p_{L|j}, \\ x_{ij,ll} = p_{ij} p_{l|j} (1 - c_{ij} p_{L|i}), \end{cases} \tag{20}$$

where $c_{ij}$ is defined by

$$\begin{aligned} 0 = &- \left(\frac{1}{i} + \frac{1}{j}\right) c_{ij} \\ &+ \frac{j-1}{j} \sum_k \frac{1}{k} \left( (1-\phi) \frac{n_{kj}}{n_j} c_{kj} (1 - c_{ji}) + \phi \frac{n\bar{d}}{ik} \frac{n_{kj} n_{ik}}{n_k n_j n_i} c_{kj} (c_{ik} - c_{ji}) \right) \\ &+ \frac{i-1}{i} \sum_k \frac{1}{k} \left( (1-\phi) \frac{n_{ki}}{n_i} c_{ki} (1 - c_{ij}) + \phi \frac{n\bar{d}}{jk} \frac{n_{ki} n_{jk}}{n_k n_i n_j} c_{ki} (c_{jk} - c_{ij}) \right). \end{aligned} \tag{21}$$

Here $\phi$ is the fraction of triangles, or triples that form a closed loop (see equation (41) in the Supplementary Material of Kuo and Carja [2024b] or equation (27) in the Supplementary Material of Kuo and Carja [2024a]). Thus, we can find $c_{ij}$ by solving this quadratic system. We then use $c_{ij}$ to express the edge dynamics and find the neutral mutant equilibrium node frequencies. Let us now analyze the mutation and selection terms and their effects on the quasi-equilibrium.

#### 1.3 Analysis of the selection term

We analyze the dynamics by considering how the quasi-equilibrium frequency $x_l$ changes when a $l$-mutant on node of degree $i$ is replaced by a $m$-mutant, from a node of degree $j$. The associated changes in $q_l$ are given by

$$\begin{aligned} m_j \to l_i : \quad \Delta x_l &= +\frac{1}{\mathbb{E}[i^{-1}]} \frac{1}{Ni} = +\Delta_i \\ l_j \to m_i : \quad \Delta x_l &= -\frac{1}{\mathbb{E}[i^{-1}]} \frac{1}{Ni} = -\Delta_i. \end{aligned} \tag{22}$$

Therefore, the amount that $x_l$ changes when a $l$-mutant on a node of degree $j$ replaces a $m$-mutant on a node of degree $i$, depends on $i$. Using equation (1), and summing across all possible node degrees $j$, we can write the total probability that $x_l$ increases by $\Delta_i$. We can similarly compute the probability that $x_l$ decreases by $\Delta_i$. The two expressions are

$$\begin{aligned} P\left(\Delta x_l = +\Delta_i\right) &= \sum_{m \neq l} \frac{(1+s)^l}{Nw} x_l x_m \sum_j n_{ij} c_{ij} \frac{1}{j} \\ P\left(\Delta x_l = -\Delta_i\right) &= \sum_{m \neq l} \frac{(1+s)^m}{Nw} x_l x_m \sum_j n_{ij} c_{ij} \frac{1}{j}. \end{aligned} \tag{23}$$

We compute the total expected change in the $x_l$, in one update and still ignoring further mutation, for now,

$$\begin{aligned} E[\Delta x_l] &= \sum_{m \neq l} \frac{(1+s)^l - (1+s)^m}{Nw} x_l x_m \frac{1}{N\mathbb{E}[i^{-1}]} \sum_{ij} n_{ij} c_{ij} \frac{1}{ij} \\ &\approx \sum_{m} \frac{(l-m)s}{N} x_l x_m \frac{1}{N\mathbb{E}[i^{-1}]} \sum_{ij} n_{ij} c_{ij} \frac{1}{ij} \end{aligned} \tag{24}$$

We only keep terms up to the linear term of $s$, and drop the $m \neq l$ condition since the $m = l$ term evaluates to 0. We further write this expected change by summing across $m$ and using $\bar{l}$ to represent the mean number of mutations in the population, which gives

$$E[\Delta x_l] = \frac{(l - \bar{l})s}{N} x_l \frac{1}{N\mathbb{E}[i^{-1}]} \sum_{ij} n_{ij} c_{ij} \frac{1}{ij}. \tag{25}$$

#### 1.4 Analysis of the mutation term

Let us now consider the effect of mutation on the quasi-equilibrium. The change in the $l$-mutant numbers due to mutation is given by

$$\begin{aligned} (l-1)_i \to l_i : \quad \Delta x_l &= +\frac{1}{\mathbb{E}[i^{-1}]} \frac{1}{Ni} = +\Delta_i \\ l_i \to (l+1)_i : \quad \Delta x_l &= -\frac{1}{\mathbb{E}[i^{-1}]} \frac{1}{Ni} = -\Delta_i. \end{aligned} \tag{26}$$

The probability that there is a mutation from the $l$-mutant into the $(l+1)$-mutant is given by

$$P(\Delta x_l = +\Delta_i \text{ due to mutation}) = \frac{U}{N} x_l. \tag{27}$$

The total expected change in the $l$-mutant numbers due to mutation is given by

$$\begin{aligned}\mathbb{E}[\Delta x_l \text{ due to mutation}] &= \frac{U}{N}(x_{l-1} - x_l)\frac{1}{\mathbb{E}[i^{-1}]}\sum_i \frac{1}{i} \\ &= \frac{U}{N}(x_{i,l-1} - x_{i,l}).\end{aligned} \tag{28}$$

#### 1.5 Adding drift

So far, to get a handle on the dynamics, we have described the dynamics of our population without drift, which stochastically affects the newly appeared mutants in the population. To incorporate stochastic effects, we need to calculate the variance and covariance in the change mutant frequency [Kimura, 1964] The variance is given by

$$\begin{aligned}Var[\Delta x_l] &\approx E[(\Delta x_l)^2] \\ &= 0\end{aligned} \tag{29}$$

Here, we only keep terms independent of $s$ and $U$. The variance is approximated by the second moment because $E[(\Delta x_l)]^2 = \mathcal{O}(s^2 + sU + U^2)$ which we drop.

Similarly, the covariance is given by

$$\begin{aligned}Cov[\Delta x_l, \Delta x_m] &\approx E[(\Delta x_l)(\Delta x_m)] \\ &= -\frac{(1+s)^l}{Nw} x_l x_m \frac{1}{N^2\mathbb{E}[i^{-1}]^2}\sum_{ij} n_{ij} c_{ij} \frac{1}{i^2 j} \\ &\approx -x_l x_m \frac{1}{N^2\mathbb{E}[i^{-1}]^2}\sum_{ij} n_{ij} c_{ij} \frac{1}{i^2 j},\end{aligned} \tag{30}$$

neglecting higher order terms of $s$ and $U$.

#### 1.6 Putting it all together and deriving the equations governing the evolutionary dynamics

Combining the above effects of the network, selection, mutation and drift terms, we now write the Fokker–Planck equation as

$$\frac{\partial P}{\partial t} = -\frac{\lambda}{2N}\sum_{l,m\neq l}\frac{\partial^2}{\partial x_l \partial x_m} x_l x_m P - \sum_l \frac{\partial}{\partial x_l}\left[\alpha\lambda s(l-\bar{l})x_l + U(x_{l-1} - x_l)\right]P, \tag{31}$$

where

$$\alpha = \mathbb{E}[i^{-1}] \sum_{ij} \frac{c_{ij} n_{ij}}{Nij} \left( \sum_{ij} \frac{c_{ij} n_{ij}}{Ni^2 j} \right)^{-1} \tag{32}$$

is the amplification factor, and

$$\lambda = \mathbb{E}[i^{-1}]^{-2} \sum_{ij} \frac{c_{ij} n_{ij}}{Ni^2 j} \tag{33}$$

is the acceleration factor. Equation (31) is the Fokker Plank equation for the probability density $P(\mathbf{x}, t|\mathbf{x}_0)$ of the multivariate random variable $\mathbf{X}_t = X_{1,t}, \cdots, X_{l,t}, \cdots, X_{b,t}$ described by the following stochastic differential equation

$$d\mathbf{X}_t = \mathbf{M}(\mathbf{X}_t)\, dt + \sqrt{\mathbf{V}(\mathbf{X}_t)}\, d\mathbf{W}_t. \tag{34}$$

Here, $\mathbf{M} = (M_1, \cdots, M_b)$, where $M_l$ is the sum of the network modified selection and mutation term for the $l$-mutant, and $\mathbf{V}$ is the covariance matrix given by equation (29) and (30). Solving for $P(\mathbf{x}, t|\mathbf{x}_0)$ gives the probability density function of the distribution of mutation numbers after time $t$, given the population starts with initial distribution of mutation numbers $\mathbf{x}_0$. This solution is our next goal.

### 2 Analysis of the equation governing the evolutionary dynamics

Fully deriving the solution to the stochastic differential equation governing mutation accumulation dynamics on a network is infeasible (or maybe just an extremely hard problem). Every node of the network has to be treated separately, and we have to consider all possible numbers of mutations on each node. This means we have to consider $Nb$ number of variables where $N$ and $b$ are arbitrarily large. However, in previous sections, we have deconstructed the stochastic differential equation governing the dynamics into parts, determined by the evolutionary forces at play, and showed that it is not necessary to model every node of the network separately. This greatly reduced the number of variables from $Nb$ to $b$.

**It is important to observe that the equation we arrive at, equation (31), is essentially that of the mutation accumulation dynamics in the well-mixed population limit [Desai and Fisher, 2007, Hallatschek, 2011], with appropriately scaled selection and drift terms, where this scaling of parameters is determined by the evolutionary properties of the network, amplification and acceleration. Therefore, if one has a a good approximation for the dynamics in the well-mixed population limit, we have shown that the approximation can be used for network structures, with the appropriate parameter stretches, as determined by the properties of the network.**

However, even this simplified equation for the well-mixed limit is very hard to solve analytically. Different approximations exist, but they all have strong assumptions on the strength of the particular evolutionary force dominating the dynamics. Therefore, they work in their respective parameter regimes, and can diverge qualitatively when their assumptions are violated. Instead of developing new analytical approaches for the well-mixed limit that fit all scenarios (beyond the scope of our study here), we rely on two previous approaches that worked well under their respective regimes.

The qualitative differences particularly appear when assuming different strengths for the force of mutation in the population, so we discuss the two mutation regimes below.

#### 2.1 Evolutionary dynamics under the weak mutation regime

For weak mutation $U \ll s$, we adopt the approach and the heuristic arguments in [Desai and Fisher, 2007]. For a detailed, rigorous derivation using generating functions and Laplace transforms, one can refer directly to [Desai and Fisher, 2007, Rouzine et al., 2008]. Here, the $x'_l$ lineages are separated into two categories and treated differently. The first category is called the bulk, where $x_l > 1/(N\alpha s\Delta l)$ and $\Delta l = l - \bar{l}$. In the bulk, the $l$-mutant frequencies is high, such that the force of selection outweighs the force of stochastic drift, and the frequencies behave deterministically [Brunet et al., 2008]. The bulk consists of the majority of the population and is concentrated near the mean of the mutation count, $\bar{l}$ [Goyal et al., 2012].

In line with these previous approaches, we define $q$ such that for $|\Delta l| > q$, the mutant is considered outside of the bulk. Therefore, we drop the stochastic component of equation (31), and write the dynamic equations as,

$$\frac{\partial x_l}{\partial t} = \alpha\lambda s(l - \bar{l})x_l + U(x_{l-1} - x_l). \tag{35}$$

Most treatments only consider the first $(q+1)$ mutants on the left of the bulk: this is because the $(q+2)-$mutant is unlikely to arise, since the $(q+1)$-mutant is in low frequency and is unable to supply $(q+1)$ with mutations. The $(q+1)$-mutant class is therefore called the lead since it is the mutant class the most fit in the population. For the lead, the mutant frequency is lower than $1/(N\alpha sq)$ and the force of drift dominates selection. Contribution from mutation depends on the previous mutant class, and can have arbitrary strength. Therefore, we must consider it. Dropping only the selection term and linearizing equation (31) in $x_q$ (since the mutant frequency is low), we write

$$\frac{\partial}{\partial t}P(x_{q+1}, t) = \frac{\lambda}{N}\frac{\partial^2}{\partial x_{q+1}^2}x_{q+1}P(x_{q+1}, t) - \frac{\partial}{\partial x_{q+1}}Ux_qP(x_{q+1}, t). \tag{36}$$

This approximation of the lead, leads to similar quantitative behavior as the heuristic argument presented by [Desai and Fisher, 2007]. By using a more sophisticated continuous time branching process to model the lead, Desai and Fisher [2007] reached the same quantitative behavior as their heuristic argument. For conciseness, we use this equation (36) for the lead dynamics.

We show in a further detailed section (**Supplementary Material Section 3.1**) that the solution to equation (35) is

$$x_l = \frac{1}{\sum_l x_l(0)e^{\alpha\lambda lst}} \sum_{m=0} x_{l-m}(0)e^{\alpha\lambda(l-m)st}\frac{r^m e^{-r}}{m!}, \quad r = \frac{U}{\alpha\lambda s}(e^{\alpha\lambda st} - 1). \tag{37}$$

As long as the contribution from a mutant class with $l > (q+1)$ is small, we replace the summation for all the mutant classes with the summation up to $q$. For weak mutation and short enough time $t \ll (\alpha\lambda s)^{-1}$, we can write

$$x_l \approx \frac{x_l(0)e^{\alpha\lambda lst}}{\sum_{l=-\infty}^{q} x_l(0)e^{\alpha\lambda lst}}. \tag{38}$$

For the frequency of the $q$-mutant, the right-most mutant class in the bulk, right before the lead, we write

$$\begin{aligned} x_q &= \frac{x_q(0)e^{\alpha\lambda qst}}{\sum_{l=-\infty}^{q} x_l(0)e^{\alpha\lambda lst}} \\ &= \frac{x_q(0)e^{\alpha\lambda qst}}{1 - x_q(0) + x_q(0)e^{\alpha\lambda qst}} \frac{1 - x_q(0) + x_q(0)e^{\alpha\lambda qst}}{\sum_{l=-\infty}^{q} x_l(0)e^{\alpha\lambda lst}} \\ &\approx \frac{x_q(0)e^{\alpha\lambda qst}}{1 - x_q(0) + x_q(0)e^{\alpha\lambda qst}}. \end{aligned} \tag{39}$$

This approximation is valid for short time $t \ll (\alpha\lambda s)^{-1}$ and also extremely long time, since the second term approaches 1 in the two limits. Using the condition $x_q(t=0) = C/N\alpha qs$, where $C$ is a constant of proportionality, that defines the transition from the bulk to the lead.

$$x_q = \frac{e^{\alpha\lambda qst}}{N\alpha qs/C - 1 + e^{\alpha\lambda qst}}. \tag{40}$$

For the dynamics at the lead, we are interested in the expected trajectory of the $(q+1)$-mutant. We get the dynamics by multiplying equation (36) by $x_{q+1}$,

$$\frac{\partial}{\partial t}\mathbb{E}[X_{q+1}, t] = \int_0^\infty x_{q+1}\frac{\lambda}{N}\frac{\partial^2}{\partial x_{q+1}^2} x_{q+1}P(x_{q+1}, t)\, dx_{q+1} - \int_0^\infty x_q \frac{\partial}{\partial x_{q+1}} U x_q P(x_{q+1}, t)\, dx_{q+1}. \tag{41}$$

Applying integration by parts on the first term on the right-hand side, we get

$$\begin{aligned}
&\int_0^\infty x_{q+1}\frac{\lambda}{N}\frac{\partial^2}{\partial x_{q+1}^2}x_{q+1}P(x_{q+1},t)\,dx_{q+1}\\
&=\left[\frac{\lambda}{N}x_{q+1}\frac{\partial}{\partial x_{q+1}}x_{q+1}P(x_{q+1},t)\right]_{x_{q+1}=0}^{x_{q+1}=\infty}-\int_0^\infty\frac{\lambda}{N}\frac{\partial}{\partial x_q}x_qP(x_q,t)\,dx_q\\
&=-\int_0^\infty\frac{\lambda}{N}\frac{\partial}{\partial x_{q+1}}x_{q+1}P(x_{q+1},t)\,dx_{q+1}\\
&=0. \qquad (42)
\end{aligned}$$

Applying integration by parts on the second term, we write

$$\begin{aligned}
-\int_0^\infty x_{q+1}\frac{\partial}{\partial x_{q+1}}Ux_qP(x_{q+1},t)\,dx_{q+1}&=x_{q+1}Ux_qP(x_{q+1},t)\Big|_0^\infty+\int_0^\infty Ux_qP(x_{q+1},t)\,dx_{q+1}\\
&=Ux_q. \qquad (43)
\end{aligned}$$

By slight abuse of notation, we define $x_{q+1}=\mathbb{E}[X_{q+1},t]$, and we essentially have

$$\frac{\partial}{\partial t}x_{q+1}=Ux_q. \qquad (44)$$

Substituting the dynamic for the $q$-mutant, we have

$$x_{q+1}(t)\approx U\int\frac{Ce^{\alpha\lambda qst}}{N\alpha qs+Ce^{\alpha\lambda qst}}\,dt.$$

We are interested in the long term stable dynamics, where dynamics is self-restoring, returning to the same state after some time. This is true when, after some time $\tau$, the mean number of mutations in the population increases by one,

$$x_{q+1}(\tau)=x_q(0)=\frac{C}{N\alpha qs}, \qquad (45)$$

and the $(q+1)$-mutant at time $\tau$ would have the same frequency as the $q$-mutant initially. At this time $\tau$, the $q$-mutant has $q$ more mutations than $\bar{l}(\tau)$, so the $q$ mutant at time 0 becomes the new $(q-1)$-mutant at time $\tau$, the $(q+1)$-mutant at time 0 becomes the $q$-mutant at time $\tau$, and so on. The dynamics would repeat itself. Based on this condition, we have

$$
\begin{aligned}
& x_{q+1}(\tau) = \frac{C}{N\alpha qs} \\
\Longrightarrow\ & NU \int_0^{\tau} \frac{Ce^{\alpha\lambda qst}\,dt}{N\alpha qs + Ce^{\alpha\lambda qst}} = \frac{C}{\alpha qs} \\
\Longrightarrow\ & \frac{NU}{\alpha\lambda qs} \int_C^{Ce^{\alpha\lambda qs\tau}} \frac{du}{N\alpha qs + u} = \frac{C}{\alpha qs} \\
\Longrightarrow\ & \Big[\ln(N\alpha qs + u)\Big]_{u=C}^{u=Ce^{\alpha\lambda qs\tau}} = \frac{C\lambda}{NU} \\
\Longrightarrow\ & [\ln(N\alpha qs + Ce^{\alpha\lambda qs\tau}) - \ln(N\alpha qs)] = \frac{C\lambda}{NU} \\
\Longrightarrow\ & \frac{N\alpha qs + Ce^{\alpha\lambda qs\tau}}{N\alpha qs + C} = \exp\left(\frac{C\lambda}{NU}\right) \\
\Longrightarrow\ & e^{\alpha\lambda qs\tau} = \frac{N\alpha qs + C}{C} \exp\left(\frac{C\lambda}{NU}\right) - \frac{N\alpha qs}{C} \\
\Longrightarrow\ & e^{\alpha\lambda qs\tau} \approx \frac{N\alpha qs}{C}\left[\exp\left(\frac{C\lambda}{NU}\right) - 1\right].
\end{aligned} \tag{46}
$$

Here, we once again use the assumption that $Ns \ll 1$. It is straightforward to get $\tau$ from the above expression.

$$
\tau = \frac{1}{\alpha\lambda qs} \ln\left\{\frac{N\alpha qs}{C}\left[\exp\left(\frac{C\lambda}{NU}\right) - 1\right]\right\}. \tag{47}
$$

This expression for $\tau$ captures the dynamics for a wide range of mutation rates $U$. For extremely weak mutation such that we are in the step-wise fixation regime, $q \approx 1$, the exponential term inside the log dominates, and we have

$$
\begin{aligned}
\tau &\approx \frac{1}{\alpha\lambda qs} \ln\left(\frac{N\alpha qs}{C}\left\{\exp\left[\frac{C\lambda}{NU}\right]\right\}\right) \\
&= \frac{1}{\alpha\lambda qs}\left(\ln\frac{N\alpha qs}{C} + \frac{C\lambda}{NU}\right) \\
&= \frac{C}{NU\alpha s} + \frac{1}{\alpha\lambda qs}\ln\frac{N\alpha qs}{C}.
\end{aligned} \tag{48}
$$

Since $\tau$ is the time for the mean mutation count to increase by one, if we let $C = 1 + \alpha s$

$$
\begin{aligned}
V &= \frac{1}{\tau} \\
&\approx \frac{NU\alpha s}{1+\alpha s},
\end{aligned} \tag{49}
$$

which is the population size times the mutation rate times the approximation of the probability of fixation. This is exactly the rate of evolution under the succession fixation regime.

To find both $\tau$ and $q$, we need another condition. We can use the fact that at time $\tau$, the mean fitness of the population would increase by 1,

$$\begin{aligned}
&\frac{\sum_l^q l x_l(0) e^{\alpha\lambda s l \tau} + (q+1)x_q(0)}{\sum_l^q x_l(0) e^{\alpha\lambda s l \tau} + x_q(0)} = 1 \\
&\implies \sum_l^q l x_l(0) e^{\alpha\lambda s l \tau} + (q+1)x_q(0) = \sum_l^q x_l(0) e^{\alpha\lambda s l \tau} + x_q(0) \\
&\implies \sum_l^q (l-1) x_l(0) e^{\alpha\lambda s l \tau} + q x_q(0) = 0 \\
&\implies x_q(0)(q-1)e^{\alpha\lambda s l \tau} = \sum_l^{q-1} (1-l) x_l(0) e^{\alpha\lambda s l \tau} - q x_q(0) \\
&\implies q - 1 = \frac{1}{x_q(0)} e^{-\alpha\lambda s q \tau} \left[ \sum_l^{q-1} (1-l) x_l(0) e^{\alpha\lambda s l \tau} - q x_q(0) \right].
\end{aligned}$$

This gives us an expression for $q$.

$$\begin{aligned}
q &= 1 + \frac{Nqs}{C} e^{-\alpha\lambda s q \tau} \left[ \sum_l^{q-1} (1-l) x_l(0) e^{\alpha\lambda s l \tau} + q x_q(0) \right] \\
&\approx 1 + \left[ \exp\left( \frac{C\lambda}{NU} \right) - 1 \right]^{-1} \left[ \sum_l^{q-1} (1-l) x_l(0) e^{\alpha\lambda s l \tau} + q x_q(0) \right].
\end{aligned}$$

We decompose the term in the second bracket into terms independent of $q$ and $\tau$, and terms that depend on them.

$$\begin{aligned}
\sum_l^{q-1} (1-l) x_l(0) e^{\alpha\lambda s l \tau} + q x_q(0) &= \sum_l^{q-1} x_l(0) e^{\alpha\lambda s l \tau} - \sum_l^{q-1} l x_l(0) e^{\alpha\lambda s l \tau} - q x_q(0) \\
&= \sum_l^{q-1} x_l(0) + \sum_l^{q-1} x_l(0) \left( e^{\alpha\lambda s l \tau} - 1 \right) - \sum_l^{q-1} l x_l(0) e^{\alpha\lambda s l \tau} - q x_q(0) \\
&= 1 + \sum_l^{q-1} x_l(0) \left( e^{\alpha\lambda s l \tau} - 1 \right) - \sum_l^{q-1} l x_l(0) e^{\alpha\lambda s l \tau} - (q+1) x_q(0) \\
&= 1 + f(N\mu).
\end{aligned}$$

We find

$$\left[\exp\left(\frac{C\lambda}{NU}\right)-1\right]^{-1} f(N\mu) \approx \frac{N\mu}{\ln(Ns/2)},$$

which gives a good fit to our simulation for a wide range of mutation, selection, and network types. Hence the approximate $q$ is given by

$$q \approx \frac{1}{1-e^{-1/NU_b}} + \frac{NU}{\ln Ns/2}. \tag{50}$$

Using equation (48) , the rate of evolution $V$ is therefore,

$$V = \frac{\alpha\lambda qs}{\ln\frac{N\alpha qs}{C} + C\lambda(NU)^{-1}} \approx \frac{q}{\lambda^{-1}\ln(N\alpha qs)+(NU)^{-1}}\frac{\alpha s}{1+\alpha s}. \tag{51}$$

#### 2.2 Evolutionary dynamics under the strong mutation regime

For large $NU$, we use the framework in Tsimring et al. [1996], Cohen et al. [2005]. Since the mutation rate is much higher than $s$, the mutant frequency at the stochastic edge, $X_{q+1}$, depends solely on the $q$-mutant supplying mutations, rather than selection. Therefore, the cutoff frequency is independent of $s$. However, the choice of $x_q$ is often arbitrary and is selected to fit simulations. For example, Cohen et al. [2005] used $x_q = 4/N$. Here, we find that the speed of evolution

$$V \approx \frac{\alpha\lambda s(\lambda^{-1}NU)^{2/3}}{\log(\alpha Ns)} + U \quad NU \gg \lambda, \tag{52}$$

folowing the $U^{2/3}$ scaling first observed by [Cohen et al., 2005, Tsimring et al., 1996]. The second term is the result of directional mutation in our model.

### 3 Additional mathematical appendix

#### 3.1 Mutation selection dynamics on graphs

In this section, we derive the solution for equation (35), which governs the dynamics in the bulk of the population. We define the generating function $G$, as

$$G(z,t;\mathbf{x}) = \sum_l e^{lz} x_l, \tag{53}$$

where, again, $\mathbf{x} = (x_1, \cdots, x_l, \cdots, x_b)$. Using the definition of $G$, we have the following relationship from equation (35),

$$\begin{aligned} \sum_l e^{lz} \frac{\partial x_l}{\partial t} &= \sum_l e^{lz} \alpha\lambda s(l - \bar{l}) x_l + \sum_l e^{lz} U(x_{l-1} - x_l) \\ \implies \frac{\partial}{\partial t} G(z,t;\mathbf{x}) &= \alpha\lambda s \left( \frac{\partial}{\partial z} - \bar{l} \right) G(z,t;\mathbf{x}) + U(e^z - 1) G(z,t;\mathbf{x}) \\ \implies \left[ \frac{\partial}{\partial t} + \alpha\lambda s \bar{l}(t) + U \right] G(z,t;\mathbf{x}) &= \left[ \alpha\lambda s \frac{\partial}{\partial z} + U e^z \right] G(z,t;\mathbf{x}). \end{aligned} \tag{54}$$

We further define

$$H(z,t;\mathbf{x}) = f(z,t) G(z,t;\mathbf{x}) = \exp\left[ \alpha\lambda s \int \bar{l}(t)\, dt + Ut + \frac{U}{\alpha\lambda s} e^z \right] G(z,t;\mathbf{x}). \tag{55}$$

This transforms equation (54) into

$$\frac{\partial}{\partial t} H(z,t;\mathbf{x}) = \alpha\lambda s \frac{\partial}{\partial z} H(z,t;\mathbf{x}). \tag{56}$$

Any function of the form $h(z + \alpha\lambda st)$ is a solution to equation (56). Using the definition of $H$ in equation (56), we express $G$ as

$$G(z,t;\mathbf{x}) = h(z + \alpha\lambda st) \exp\left[ -\alpha\lambda s \int \bar{l}(t)\, dt - Ut - \frac{U}{\alpha\lambda s}(e^z - 1) \right]. \tag{57}$$

We can explicitly solve for $h$ utilizing the two boundary conditions:

$$G(z,0;\mathbf{x}) = \sum_l x_l(0) e^{lz}, \tag{58}$$

and

$$G(0,t;\mathbf{x}) = \sum_l x_l(t) = 1. \tag{59}$$

Let $t = 0$, and arbitrarily set $l(0) = 0$, meaning the mean mutation count in the population is initially 0 at time $t = 0$. With this definition, we have

$$\begin{aligned} G(z,0;\mathbf{x}) &= h(z)\exp\left[-\frac{U}{\alpha\lambda s}(e^z - 1)\right] \\ \implies h(z) &= G(z,0;\mathbf{x})\exp\left[\frac{U}{\alpha\lambda s}(e^z - 1)\right]. \end{aligned} \tag{60}$$

We can easily get

$$h(z+st) = G(z+\alpha\lambda st,0;\mathbf{x})\exp\left[\frac{U}{\alpha\lambda s}(e^{z+\alpha\lambda st} - 1)\right], \tag{61}$$

and thus,

$$\begin{aligned} G(z,t;\mathbf{x}) &= G(z+\alpha\lambda st,0;\mathbf{x})\exp\left[\frac{U}{\alpha\lambda s}(e^{z+\alpha\lambda st} - 1)\right]\exp\left[-\alpha\lambda s\int \bar{l}(t)\,dt - Ut - \frac{U}{\alpha\lambda s}(e^z - 1)\right] \\ &= G(z+\alpha\lambda st,0;\mathbf{x})\exp\left[\frac{U}{\alpha\lambda s}e^z(e^{\alpha\lambda st} - 1) - \alpha\lambda s\int \bar{l}(t)\,dt - Ut\right] \\ &= G(z+\alpha\lambda st,0;\mathbf{x})\exp\left[\frac{U}{\alpha\lambda s}e^z(e^{\alpha\lambda st} - 1)\right]\exp\left[-\alpha\lambda s\int \bar{l}(t)\,dt - Ut\right]. \end{aligned} \tag{62}$$

Let $z = 0$,

$$\begin{aligned} 1 &= G(\alpha\lambda st,0;\mathbf{x})\exp\left[\frac{U}{\alpha\lambda s}(e^{\alpha\lambda st} - 1)\right]\exp\left[-\alpha\lambda s\int \bar{l}(t)\,dt - Ut\right] \\ \exp\left[\alpha\lambda s\int \bar{l}(t)\,dt + Ut\right] &= G(\alpha\lambda st,0;\mathbf{x})\exp\left[\frac{U}{\alpha\lambda s}(e^{\alpha\lambda st} - 1)\right]. \end{aligned} \tag{63}$$

Substitute this value into $G$

$$\begin{aligned} G(z,t;\mathbf{x}) &= \frac{G(z+\alpha\lambda st,0;\mathbf{x})}{G(st,0;\mathbf{x})}\exp\left[\frac{U}{\alpha\lambda s}(e^z - 1)(e^{\alpha\lambda st} - 1)\right] \\ &= \frac{\sum_l x_l(0)e^{lz+\alpha\lambda ls}}{\sum_l x_l(0)e^{\alpha\lambda lst}}\exp\left[\frac{U}{\alpha\lambda s}(e^z - 1)(e^{\alpha\lambda st} - 1)\right]. \end{aligned} \tag{64}$$

The first term

$$G_1(z,t;\mathbf{x}) = \frac{\sum_l x_l(0)e^{lz+\alpha\lambda lst}}{\sum_l x_l(0)e^{\alpha\lambda lst}}, \tag{65}$$

is the generating function of

$$x_l = \frac{x_l(0)e^{\alpha\lambda lst}}{\sum_l x_l(0)e^{\alpha\lambda lst}}. \tag{66}$$

The second term

$$G_2(z,t;\mathbf{x}) = \exp\left[\frac{U}{\alpha\lambda s}(e^z-1)(e^{\alpha\lambda st}-1)\right]. \tag{67}$$

is the moment generating function for Poisson distribution. This means

$$x_{l,2} = \frac{r^l e^{-r}}{l!}, \quad r = \frac{U}{\alpha\lambda s}(e^{\alpha\lambda st}-1). \tag{68}$$

Using the additive property of generating functions, we obtain

$$x_l = \sum_{m=0} \frac{x_{l-m}(0)e^{\alpha\lambda(l-m)st}}{\sum_l x_l(0)e^{\alpha\lambda lst}} \frac{r^m e^{-r}}{m!}. \tag{69}$$

#### 3.2 Deriving the amplification and acceleration factors for regular graphs and bipartite graphs

In **Supplementary Material section 1**, we showed the how to derive the network amplification and acceleration factors analytically (equations 32 and 33). This involves solving equation (21). This has to be done numerically, if the network has nodes of many distinct degrees. However, we demonstrate that when the graph contains only one or two distinct degrees, equation (21) yields straightforward closed-form approximations for the network amplification and deceleration factors, such as in the case of regular and bipartite graphs.

##### 3.2.1 Analytics for the evolutionary properties of regular graphs

A regular graph is a graph where each node has the same number of neighbors; i.e. every node has the same degree $i$. This simplifies equation (21) to

$$i = (i-1)(1-\phi)\frac{n_{ii}}{n_i}(1-c_{ii}). \tag{70}$$

In a regular graph, the number of $ii$ edges is simply the number of $n_i$ nodes times $i$, $n_{ii} = in_i$. Thus,

$$c_{ii} = 1 - \frac{1}{(i-1)(1-\phi)}. \tag{71}$$

Using the approximation of the amplification factor in equation (32),

$$\begin{aligned} \alpha &= E[i^{-1}] \sum_{ij} \frac{c_{ij} n_{ij}}{Nij} \left( \sum_{ij} \frac{c_{ij} n_{ij}}{Ni^2 j} \right)^{-1} \\ &= i^{-1} \frac{c_{ii} n_{ii}}{Ni^2} \left( \frac{c_{ii} n_{ii}}{Ni^3} \right)^{-1} \\ &= 1. \end{aligned} \tag{72}$$

Using the approximation of the deceleration factor in equation (33)

$$\begin{aligned} \lambda &= E[i^{-1}]^{-2} \sum_{ij} \frac{c_{ij} n_{ij}}{Ni^2 j} \\ &= i^2 \frac{c_{ii} n_{ii}}{Ni^3} \\ &= c_{ii}. \end{aligned} \tag{73}$$

##### 3.2.2 Analytics for the evolutionary properties of bipartite graphs

A bipartite graph is a graph whose nodes can be divided into two node groups: one with degree $i$ and one with degree $j$. All the edges in the graph are of type $ij$, that is every edge connects a node of degree i and a node of degree $j$. Every node with degree $i$ is connected to all nodes of degree $j$, and vice versa. As a result, $c_{ii}$ and $c_{jj}$ are irrelevant in equation (21). There are also no triangles in bipartite graphs, since there is no $ii$ and $jj$ edges to close the loop. Therefore equation (21) becomes

$$\begin{aligned} &\left( \frac{1}{i} + \frac{1}{j} \right) = \frac{j-1}{ij} \frac{n_{ij}}{n_j} (1 - c_{ij}) + \frac{i-1}{ij} \frac{n_{ji}}{n_i} (1 - c) \\ \implies\ & i + j = (j-1) \frac{n_{ij}}{n_j} (1 - c_{ij}) + (i-1) \frac{n_{ji}}{n_i} (1 - c_{ij}). \end{aligned} \tag{74}$$

Every node with degree $i$ is connected to all nodes of degree $j$, and vice versa $n_i = j, n_j = i, n_{ij} = ij$, and $i + j = N$. This lets us express $c$ in terms of the population size, $N$, and node degrees $i$ and $j$,

$$\begin{aligned}
N &= j(j-1)(1-c_{ij}) + i(i-1)(1-c_{ij}) \\
&= (N-i)(N-i-1)(1-c_{ij}) + i(i-1)(1-c_{ij}) \\
&= (N^2 - Ni - N - Ni + i^2 + i)(1-c_{ij}) + i(i-1)(1-c_{ij}) \\
&= (N^2 - 2Ni - N + 2i^2)(1-c_{ij}) \\
c_{ij} &= 1 - \frac{N}{N^2 - 2Ni - N + 2i^2}. \qquad (75)
\end{aligned}$$

Using the approximation of the amplification factor in equation (32),

$$\begin{aligned}
\alpha &= E[i^{-1}] \sum_{ij} \frac{c_{ij} n_{ij}}{Nij} \left( \sum_{ij} \frac{c_{ij} n_{ij}}{Ni^2 j} \right)^{-1} \\
&= \frac{1}{N} \left( \frac{i}{j} + \frac{j}{i} \right) 2 \frac{cij}{Nij} \left( \frac{cij}{Ni^2 j} + \frac{cij}{Nij^2} \right)^{-1} \\
&= 2 \frac{i^2 + j^2}{Nij} \left( \frac{1}{i} + \frac{1}{j} \right)^{-1} \\
&= 2 \frac{i^2 + j^2}{N(i+j)} \\
&= 2 \frac{i^2 + j^2}{N^2}. \qquad (76)
\end{aligned}$$

Using the approximation of the deceleration factor in equation (33)

$$\begin{aligned}
\lambda &= E[i^{-1}]^{-2} \sum_{ij} \frac{c_{ij} n_{ij}}{Ni^2 j} \qquad (77) \\
&= \frac{1}{N^2} \left( \frac{i}{j} + \frac{j}{i} \right)^{-2} \left( \frac{c_{ij} ij}{Ni^2 j} + \frac{c_{ij} ij}{Nij^2} \right) \\
&= \frac{1}{N^2} \left( \frac{i^2 + j^2}{ij} \right)^{-2} \frac{c_{ij} ij}{Nij} \left( \frac{1}{i} + \frac{1}{j} \right) \\
&= \frac{1}{N^2} \left( \frac{i^2 + j^2}{ij} \right)^{-2} \frac{c_{ij} ij}{Nij} \frac{N}{ij} \\
&= \frac{cijN^2}{(i^2 + j^2)^2}. \qquad (78)
\end{aligned}$$

For large bipartite graphs, $c_{ij}$ is approximately 1, thus

$$\lambda \approx \frac{ijN^2}{(i^2+j^2)^2}. \tag{79}$$

We also get the following relationship between the deceleration factor and amplification factor.

$$\begin{aligned}\lambda &= \frac{4i(N-i)}{N^2\alpha^2} \\ &= \frac{2-\alpha}{\alpha^2} \\ &= \frac{2}{\alpha^2} - \frac{1}{\alpha}.\end{aligned} \tag{80}$$

### 4 Supplementary Figures

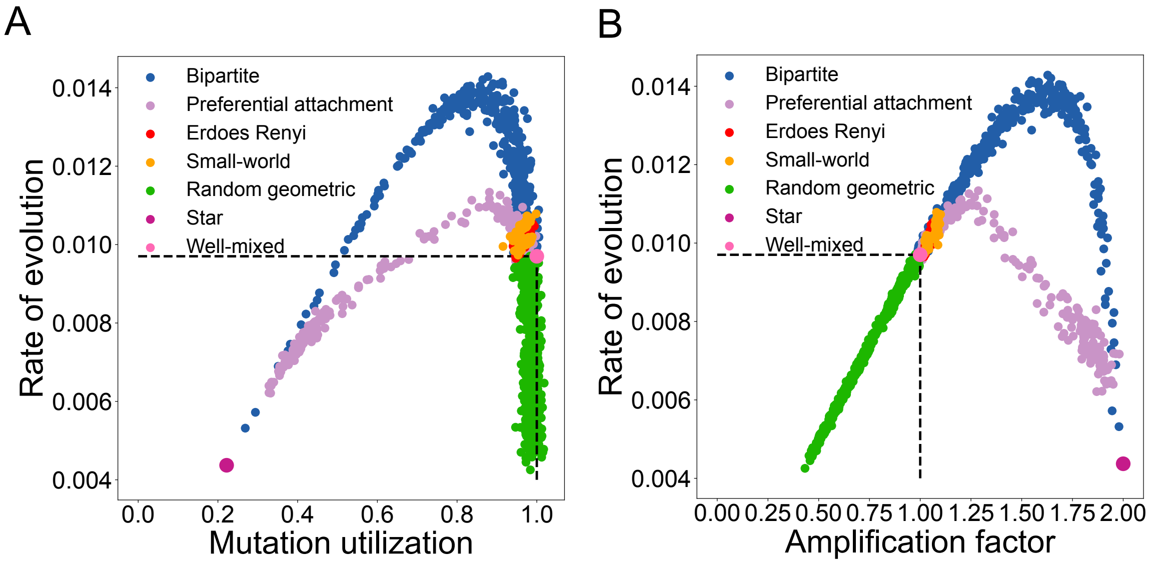

Supplementary Figure S1: **The rate of evolution and genetic variation across network families under weak mutation. Panel A.** Here, the mutation rate is set to $10^{-5}$, $N = 1000$ and $Ns = 10$. The dots represent individual networks of the families depicted. Mutation utilization is calculated by taking the rate of evolution divided by the probability of fixation of a mutant on the network (rate of evolution in the limit of no mutation). **Panel B.** The rate of evolution, as a function of the amplification and acceleration factor for graphs of size 1000. The mutation rate is set to $10^{-5}$ and, $Ns = 10$. The dots represent individual networks of the families depicted.

### References

Éric Brunet, Igor M Rouzine, and Claus O Wilke. The stochastic edge in adaptive evolution. *Genetics*, 179 (1):603–620, 2008.

Elisheva Cohen, David A Kessler, and Herbert Levine. Front propagation up a reaction rate gradient. *Physical Review E*, 72(6):066126, 2005.

Michael M Desai and Daniel S Fisher. Beneficial mutation–selection balance and the effect of linkage on positive selection. *Genetics*, 176(3):1759–1798, 2007.

Sidhartha Goyal, Daniel J Balick, Elizabeth R Jerison, Richard A Neher, Boris I Shraiman, and Michael M Desai. Dynamic mutation–selection balance as an evolutionary attractor. *Genetics*, 191(4):1309–1319, 2012.

Oskar Hallatschek. The noisy edge of traveling waves. *Proceedings of the National Academy of Sciences*, 108 (5):1783–1787, 2011.

Motoo Kimura. Diffusion models in population genetics. *Journal of Applied Probability*, 1(2):177–232, 1964.

Motoo Kimura. The number of heterozygous nucleotide sites maintained in a finite population due to steady flux of mutations. *Genetics*, 61(4):893, 1969.

Yang Ping Kuo and Oana Carja. Evolutionary graph theory beyond single mutation dynamics: on how network structured populations cross fitness landscapes. *Genetics*, page iyae055, 2024a.

Yang Ping Kuo and Oana Carja. Evolutionary graph theory beyond pairwise interactions: Higher-order network motifs shape times to fixation in structured populations. *PLOS Computational Biology*, 20(3): e1011905, 2024b.

Emanuele Pugliese and Claudio Castellano. Heterogeneous pair approximation for voter models on networks. *Europhysics Letters*, 88(5):58004, 2009.

Igor M Rouzine, Éric Brunet, and Claus O Wilke. The traveling-wave approach to asexual evolution: Muller ratchet and speed of adaptation. *Theoretical Population Biology*, 73(1):24–46, 2008.

Lev S Tsimring, Herbert Levine, and David A Kessler. RNA virus evolution via a fitness-space model. *Physical Review Letters*, 76(23):4440, 1996.
